# Loss of direct binding of *Leishmania* profilin with actin adversely affected its functions and interactions with other cellular proteins

**DOI:** 10.1101/2025.04.17.649292

**Authors:** Srikalaivani Raja, Prashant K. Rai, S. Hemant Kumar, S. Thiyagarajan, Chhitar M. Gupta

## Abstract

Profilins are actin binding proteins that play a central role in regulation of actin remodeling in all eukaryotic organisms. *Leishmania,* a family of protozoan parasites that cause leishmaniasis, express a single homolog of profilin, which differs from other eukaryotic profilins in that it contains an extra stretch of 20 amino acids (actin binding domain) through which it directly binds actin. We therefore considered it of interest to analyze the role of direct binding of profilin with actin in interactions and functions of profilin in *L. donovani* promastigotes. For this, we deleted the actin binding domain in *L. donovani* profilin (LdPfn), and then carried out comparative analysis of LdPfn and truncated LdPfn (δLdPfn) interactomes by affinity pull-down and mass spectrometry. To assess the effect of deletion of the actin binding domain on the LdPfn functions, we expressed GFP conjugates of LdPfn and δLdPfn in the wild type *Leishmania* cells and then carried out comparative analysis of their growth, cell division cycle and intracellular vesicle trafficking activity. Results revealed that expression of GFP-δLdPfn in the wild type cells adversely affected their cell growth, intracellular vesicle trafficking and G1-to-S and S-to-G2/M phase transitions during their cell division cycle. Also, there was a complete loss of LdPfn binding to several cellular proteins including actin and mitochondrial outer membrane protein, porin (LdPorin) after deleting its actin binding domain. Further, to assess the effect of lack of LdPorin binding to δLdPfn on the mitochondrial functions, we measured the cellular ATP levels in wild-type and transgenic promastigotes. The ATP levels were significantly increased by expressing GFP-δLdPfn in wild-type cells. These results taken together strongly indicate that LdPfn-driven actin remodeling besides playing a pivotal role in regulation of intracellular vesicle trafficking and cell division cycle, it also plays an important role in regulation of *Leishmania* mitochondrial activity.

## Introduction

Profilins are multi-ligand binding proteins which play a central role in regulation of actin dynamics in most eukaryotic cells [1]. On one hand, they catalyze ADP to ATP exchange on ADP-actin and thereby promote actin polymerization process, while on the other hand, they sequester actin to maintain optimum concentration of actin monomers within the cytoplasm [1]. In addition to actin, profilins also bind to Poly-L-Proline (PLP) stretches and polyphosphoinositides [2]. Profilin -driven actin remodeling is involved in several vital cellular processes, such as motility, endocytosis, intracellular vesicle trafficking, etc. [1,2]. Besides being present in higher eukaryotic cells, profilin is also present in lower eukaryotic organisms, including trypanosomatid parasites, such as *Leishmania* [3].

*Leishmania* is a family of flagellated protozoan parasites that cause several human diseases, including visceral leishmaniasis, also known as kala-azar, which if left untreated becomes fatal in over 90% cases [4]. These parasites exist in two forms, which are morphologically quite distinct. While the oval-shaped and non-motile amastigotes reside and multiply within the host mammalian macrophages, the spindle-shaped and highly motile promastigotes grow and multiply within the alimentary canal of the sand fly vector [5]. *Leishmania*, similarly to other eukaryotic organisms, contain genes that encode actin and a limited repertoire of actin binding and actin related proteins [6].

Earlier studies with *L. donovani* promastigotes revealed that these parasites express a single homolog of profilin (LdPfn) that localizes throughout the cell body, including the flagellum, nucleus and the kinetoplast [3]. Similarly to other eukaryotic profilins, LdPfn at low concentration promotes actin polymerization by catalyzing the exchange of ADP for ATP on ADP-bound actin, but at high concentration, it sequesters actin monomers [3]. It has further been shown that LdPfn is involved in regulation of intracellular vesicle trafficking [3] and cell division cycle [7]. As *Leishmania* profilins, unlike other profilins, contain an extra stretch of 20 amino acids in their amino acid sequences through which *L. major* profilin (LmPfn) has been reported to bind to *L. major* actin (LmAct) [8], we thought it appropriate to investigate the role of direct binding of profilin with actin in its functions and interactions with other cellular proteins in *L. donovani* promastigotes. For this, we deleted the extra stretch of 20 amino acids from the LdPfn amino acids sequence and then analyzed its effects on the LdPfn interactions and functions.

## Materials and Methods

### Leishmania culture

*L. donovani* (Strain: MHOM/IN/80/DD8; ATCC Cat.No.50212) cells were maintained in high glucose Dulbecco’s modified Eagle’s medium (DMEM) (Life Technologies, Thermo-Fisher Scientific) supplemented with 10% heat inactivated fetal bovine serum (MP chemicals) and 4 mg/ml gentamicin (MP chemicals).

### Cloning of LdPfn and amino acids 110 to 128 deleted LdPfn (**δ**LdPfn) genes containing glutathione-S-transferase (GST) or green fluorescence protein (GFP) tag at their N-terminal

LdPfn gene construct with GST tag at its N-terminal (GST-LdPfn) was cloned in pGEX-4T2 vector, whereas LdPfn gene construct with GFP tag at its N-terminal was cloned in pXG-GFP^+2^ vector, as described earlier [3]. For preparing the gene constructs of GST-δLdPfn and GFP-δLdPfn, the gene constructs of GST-LdPfn and GFP-LdPfn were used as the templates, which were first PCR amplified, using primers listed in S1Table, by subjecting them into initial denaturation at 95°C for 5 min and then 15 cycles of denaturation at 95°C for 30 sec. This was followed by annealing (63.3°C for 1 min) and extension (72°C for 10 min). The final extension step was at 72°C for 30 min. Phusion DNA polymerase was used in all reactions. Q5 ® Site-Directed Mutagenesis Kit was used for selecting the deletion mutants. The reaction mixture contained PCR product (1μl), Kinase, Ligase and Dpn1 (KLD) reaction buffer (5ul), KLD enzyme mixture (1μl) and nuclease free water (3ul) mixed well by pipetting up and down at room temperature for 5 min. DH5α competent cells were transformed using 5 μl of treated PCR products, plated on ampicillin Luria-Bertani (LB) agar plates and incubated at 37°C. The positive clones were confirmed by restriction digestion followed by Sanger sequencing.

### Expression and purification of GST-LdPfn and GST-δLdPfn proteins

GST-LdPfn and GST-δLdPfn were expressed in *Escherichia coli* BL21 (DE3), as reported earlier [3]. Proteins were purified, using an AKTA chromatography system (GE Healthcare) on a Superdex 200 column. Proteins were eluted, using 20mM Tris-HCl buffer, pH 7.5, containing 300mM sodium chloride, 5% glycerol and 1mM EDTA as 2 ml fractions (S2 Fig). The pure fractions were pooled together, concentrated, and then stored at -80°C

### Cloning and expression of gene encoding mitochondrial outer membrane protein, porin, containing glutathione-S-transferase (GST) tag at its N-terminal and protein purification

Mitochondrial outer membrane protein, porin, putative Gene ID LDBPK_020430.1, (LdPorin) was amplified by PCR, using *L donovani* genomic DNA as the template. The forward and reverse primers used are listed in S2 Table. The PCR amplified product (876 base pairs) was cloned in pGEX-4T2 vector at cloning sites Bam HI and Xho1 in frame with N-terminal GST tag. LdPorin was expressed in *Escherichia coli* BL21 p-lysis. Transformed cells were grown at 37 °C to an OD600 of 0.6 in LB media and the expression was induced with 0.5 mM isopropyl β-D-thiogalactoside (IPTG, MP Biochemicals) at 30 °C for 6h. Bacterial cells were pelleted and re-suspended in buffer containing 10 mM Tris-HCl, pH 7, 150 mM NaCl, 0.1 mM EDTA, 1% Triton X-100, 1 mM DTT, 30% sucrose and lysed by mild sonication on ice. Lysates were cleared by centrifugation at 12000×g for 30 min at 4°C and the supernatant was passed through a pre-equilibrated GSTrap™ 4B column (GE Healthcare). Column was washed with PBS, 0.1% Triton X-100, 1 mM DTT and bound protein eluted with 50 mM glutathione in 50 mM Tris-HCl, pH 8.0. Similarly, pGEX-4T2 vector alone was expressed and the protein was purified using the same protocol. Eluate was concentrated and resolved on SDS-polyacrylamide gel by electrophoresis (S3 Fig)

### Affinity Pull-down and mass spectrometry

Comparative analysis of LdPfn and δLdPfn interactomes was carried out using affinity pull-down-mass spectrometry assay, essentially following our published procedure [7]. Briefly, purified GST-LdPfn, GST δLdPfn and GST alone were first incubated with Glutathione-Sepharose affinity beads. The beads were washed to remove unbound proteins and then the bead bound recombinant proteins were incubated with clear *L. donovani* promastigotes lysates. The unbound proteins were removed by washing, and from the washed beads, the *Leishmania* proteins were eluted by boiling the beads in SDS-PAGE buffer at 96^0^ C for 5 min. The eluted protein samples were resolved on SDS-polyacrylamide gel (12%) by electrophoresis. Three biological replicates of each of GST-LdPfn, GST-δLdPfn and GST alone pull-down samples were rinsed three times with double distilled water; the respective lanes were excised and submitted at the mass spectrometry facility (Centre for Cellular and Molecular platforms, GKVK Campus, Bellary Road, Bangalore Karnataka, India) for analysis, using LC-MS ((Orbitrap Fusion™ Tribrid™ Mass Spectrometer). Only the protein hits with a score cut off >40 and significance threshold p<0.05 were considered. The data thus-obtained were used for searching the www.tritrypdb.org database (version 51), using the *L. donovani* (BPK282A1) strain as reference. Data has been deposited at PRIDE–Proteome X-change Consortium [9] and may be accessed with the identifier: PXD039942. Among the three biological replicates, the proteins present in eluates of all the three replicates of GST-LdPfn pull down, but not present in eluates of any of the three replicates of GST-δLdPfn or GST alone pull down, were taken into consideration.

### Poly -L-proline (PLP) binding

Equal amounts of purified GST-LdPfn and GST-δLdPfn proteins (1mg/ml) were allowed to bind with PLP-coupled Sepharose 4B columns for 1 h. The columns were washed with 4 M urea, 10 mM Tris, pH 8.0, 1 mM EDTA, 100 mM NaCl and 2 mM DTT. Column-bound GST-LdPfn and GST-δLdPfn were eluted with 8 M urea buffer (8 M urea, 10 mM Tris pH 8.0, 1 mM EDTA, 100 mM NaCl and 2 mM DTT). The PLP-coupled Sepharose 4B resin was prepared by coupling poly-L-proline having a molecular weight 1000−10000 Da (Sigma Aldrich, Cat.no P2254) with cyanogen bromide-activated-Sepharose 4B resin, following the manufacturer’s protocol.

### Western blotting

*Leishmania* cell lysates were prepared by re-suspending and boiling the cell pellets for 5 min in SDS sample buffer. Samples were resolved on SDS-polyacrylamide gel by electrophoresis and electro-blotted onto nitrocellulose membrane (0.45μm, Synergy scientific services) in Tris - glycine buffer (pH 8.3) at 100V for 2 h. The membrane was treated with 5% skimmed milk to block the nonspecific sites, and then probed with rabbit anti-LdPfn antibodies (1:100) or chicken anti-*L. donovani*actin (LdAct) antibodies (1:5000) for overnight at 4^0^C. The membrane was washed 5 times with Tris-buffered saline (pH 7.5) containing 0.05% (v/v) Tween 20 (TBST), then incubated with HRP conjugated anti-rabbit IgG antibody (Invitrogen, cat no. A16074) or HRP conjugated anti-chicken IgY antibody (Invitrogen) for 2 h. After 2h incubation, it was washed three times with TBST buffer at 15 min intervals and developed with ECL (Bio-Rad, Clarity) and imaging software (Syngene, G-box).

### ATP estimation

ATP standards were prepared by diluting the 10 mM ATP solution to final concentrations of 0 (blank), 200, 400, 600, 800, and 1000 picomoles/well. ATP assay buffer was added to each well followed by the ATP probe, ATP converter, and developer mix according to the manufacturer’s protocol to make-up the volume to 100 µl. The reaction mixture was incubated for 30 min at room temperature. Finally, fluorescence was read at λ_ex_ = 535nm / λ_em_ = 587nm on a microplate reader. The R^2^ value (0.9923) was calculated from the standard curve.

For cell sample preparation, 1× 10^6^ cells were lysed in 100 µl of ATP assay buffer and then deproteinized using a 10 kDa MWCO spin filter. 50 µl of each sample was added to the well followed by the ATP assay buffer, ATP probe, ATP converter, and developer (as per manufacturer protocol) to make-up final volume to 100µl, and mixed by pipetting. Fluorescence was read at λ_ex_ = 535nm / λ_em_ = 587nm on a microplate reader. The amounts of ATP in cell samples were then calculated from the ATP standard curve

### Bioinformatics analysis

Actin-proflin complex structures of *Homo sapiens* (Hs)and *Leishmania major* (Lm) were downloaded from www.rcsb.org (PDB IDs 6nbw and 8c47, respectively), and that of *Leishmania donovani* (Ld) was predicted using the AlphaFold server [10]. Profilin structures of *Homo sapiens, Leishmania major* and *Leishmania donovani* were culled from their respective actin-profilin complexes, while those of *Leishmania infantum* and *Leishmania mexicana* were downloaded from AlphaFold [11–13]. Human N-WASP structure was obtained from www.rcsb.org(PDB ID 2vcp) and the structure of WH2 domain of LdPfn was culled from the AlphaFold generated structure of the same. Structure based multiple sequence alignment for profilins was performed using the Dali server [14] while that of the human N-WASP and WH2 domain of LdPfn by using the software suite ICM.

### Cell growth analysis

For cell growth analysis, the wild type cells were transfected separately with GFP-LdPfn and GFP-δLdPfn gene constructs. The transfected cells were grown in DMEM containing 10% fetal bovine serum, without antibiotics. Initial cell density was 5 × 10^5^ cells /ml and the cell growth was recorded at every 24 h interval, using a hemocytometer.

### Intracellular trafficking

The lipid soluble fluorescent dye FM™4−64 FX was used for studying the intracellular vesicle trafficking in *Leishmania* cells, as described by Sahin et al. [15]. Briefly, the cells (10^7^ /ml) were incubated in culture medium containing 10% fetal calf serum and 2 μg/ml FM4-64 FX (Invitrogen) for 10 min at 25°C in dark. The cells were harvested, resuspended in fresh medium, and incubated for varying time intervals at 25^0^C. After two washings with cold PBS solution, the cells collected at various time points were spread onto cover slips, fixed with 2% paraformaldehyde, washed, mounted and images were captured on a fluorescence microscope (Nikon laser scanning confocal microscope c2).

### Cell division cycle

Analysis of cell division cycle of GFP-LdPfn and GFP-δLdPfn expressing cells was carried out using flow cytometry. For this, *Leishmania* cultures (5 x 10^7^ cells) were synchronized by incubating them with 200 mg /ml of N-hydroxy urea (HU) (Sigma) overnight (12–14 h). The cells were washed and then re-suspended in DMEM media containing 10% fetal calf serum without HU. Small aliquots of cell suspension were withdrawn at 2h interval up to 10 h. The cells after washing, were resuspended in 50μl of PBS and mixed with 150μl of fixative solution (1% Triton X-100, 40mM citric acid, 20mM sodium phosphate, 200mM sucrose). The mixture was incubated at 25^0^ C for 5 min. After adding 350 μl of diluent buffer (125mM MgCl_2_ in PBS), the cells were stored at 4^0^ C until further use. Before the cell cycle analysis, the fixed cells were treated with 50μg RNase (5mg / ml in 0.2M sodium phosphate buffer, pH7.0) for 2 h at 37°^0^ C and then incubated with 50μg PI (5mg /ml in 1.12% sodium citrate) for 30 min at 25^0^ C. The samples were analyzed in Gallios flow cytometer (Beckman coulter) and proportions of the G1, S, and G2M populations were determined using Mod Fit LT software (Verity Software House, Topsham, ME, USA). All the experiments were done in triplicates.

### Statistical Analysis

All the experiments were carried out at least three times and the results were expressed as the standard deviation (SD) of the mean (mean ± SD) of three experiments. The data were statistically analyzed by using the Student’s t-test. A p-value of <0.05 was considered significant.

## Results

*Leishmania* profilins differ from *Homo sapiens* profilins (HsPfn) in that they contain an extra stretch of about 20 amino acids (aa) in their aa sequence (Fig.1A), which is present as an α-helical insertion in their 3-d structures (Fig.1B). To delineate the differences between the structures of *Leishmania* and *Homo sapiens* profilins, we superimposed the structures of *L. major* profilin (LmPfn), *L. donovani* profilin (LdPfn) and HsPfn on each other (Fig.1B). Results revealed that although overall folds in all the three profilins structures are largely conserved, difference between these two classes profilins lies mainly because of α-helical insertion present in structures of *Leishmania* profilins (Fig.1B). The α-helical insertion exhibited high similarity with the Wiskott-Aldrich syndrome (WAS) protein homology 2 (WH2) domain (Fig.1C), found in several actin monomer binding proteins [16]. Further, it has recently been reported that LmPfn interacts with Lm Actin (LmAct) through the WH2 -like domain [8]. To ascertain whether LdPfn interacts with Ld Actin (LdAct) with a similar mechanism, we overlaid the structure of the LmPfn-LmAct complex on the modeled structure of LdPfn-LdAct complex. Results given in Fig.2A indicate that LdPfn interacts with LdAct in almost identical way, compared to LmPfn-LmAct interaction. To show that *Leishmania* profilin – actin interactions are different from that of HsPfn-HsAct interaction, we superimposed the structures of HsPfn-HsAct, LmPfn-LmAct and LdPfn-LdAct complexes on each other. Fig.2B shows that the mechanism involved in direct interaction of profilin with actin in *Leishmania* is significantly different from that of other eukaryotic organisms.

**Fig 1:**
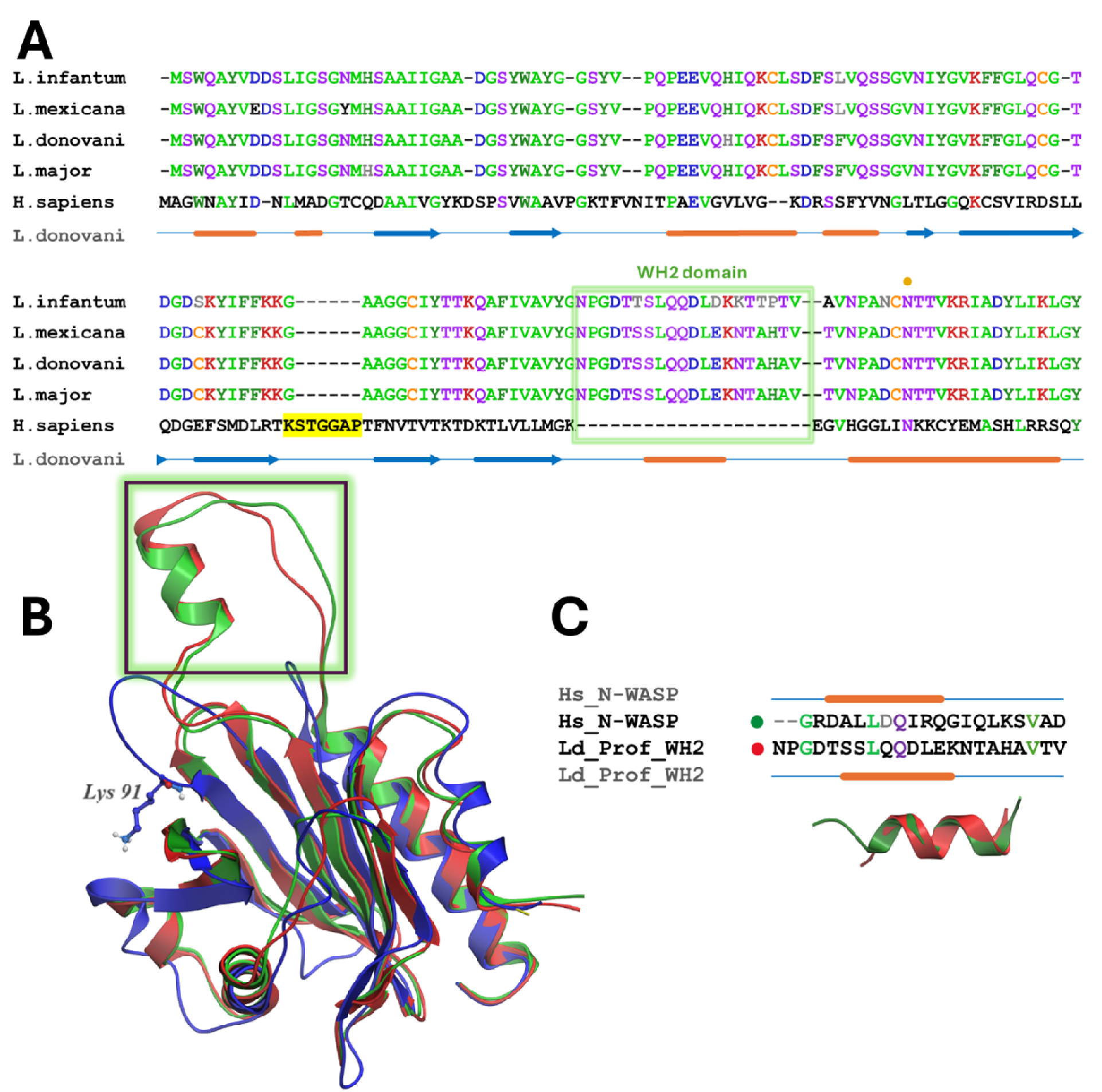
**A) Structure-based sequence alignment of profilin in *L. infantum, L. mexicana, L. donovani, L major* and *Homo sapiens***. WH2 domain is depicted in a green box. Lys91 containing loop of HsPfn is highlighted in yellow. Conserved Asn125 is denotated by a golden dot above the sequence block. Secondary structure regions as inferred from *L. donovani* structure are marked as lines (helices) and arrows (strands) below the sequence block. **B)** Superimposition of LdPfn (red), LmPfn (green) and HsPfn (blue). WH2 domain is marked **by a black rectangle**. LdPfn structure was culled from actin-profilin complexes generated using the AlphaFold server. LmPfn and HsPfn structures were culled from their respective crystal structures (PDB IDs 8c47 and 6nbw); Lys91 shown in sticks indicate the additional loop in human profilin that interacts with actin. **C) Sequence alignment of WH2 domain in LdPfn and N-WASP of humans** (PDB ID 2vcf), and **overlay of the helices in the human N-WASP (green) and WH2 domain of LdPfn (red).** LdPfn, *Leishmania donovani* profilin.

**Fig 2:**
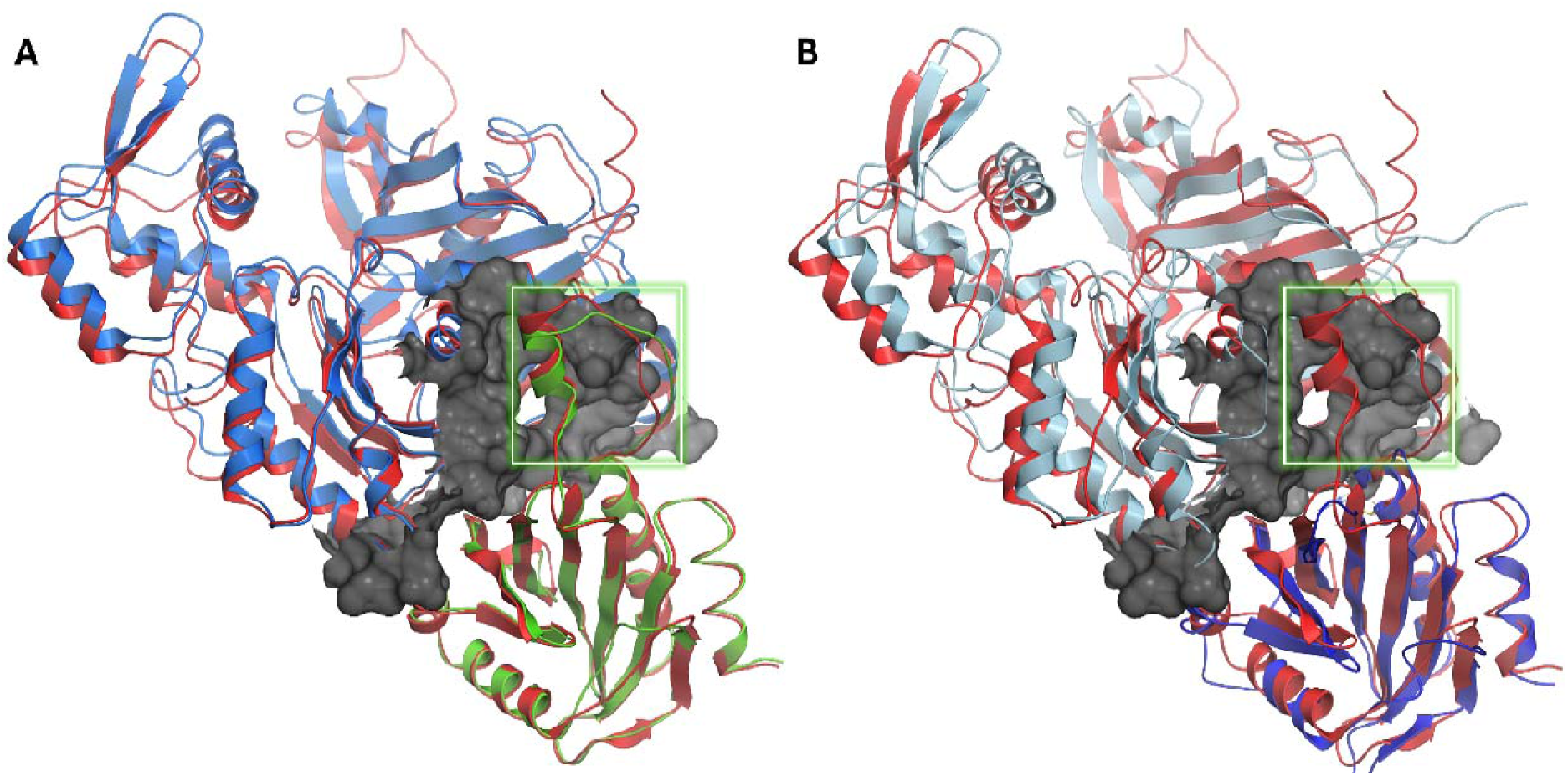
**A) Overlay of actin-profilin complexes of *L. donovani* (red) and *L major* (blue-green).** LdPfn-LdAct complex was predicted using AlphaFold server while LmPfn-LmAct complex was obtained from PDB (ID 8c47). **B) Overlay of actin-profilin complexes of *L. donovani* (red)and *H. sapiens* (cyan-blue).** Structure of LdPfn-LdAct complex was predicted using AlphaFold server while HsPfn-HsAct complex was obtained from PDB (ID 6nbw). WH2 domains are highlighted in a green box.

It is interesting to observe that the mode of direct binding of profilin to actin in humans and *Leishmania* is entirely different. We therefore thought it appropriate to investigate the role of direct binding of LdPfn to LdAct in LdPfn functions and interactions with other cellular proteins to assess the suitability of LdPfn-LdAct interaction as a drug target. Keeping this in view, we deleted the helical insert region (aa110 to aa128; total 19 aa) from the LdPfn aa sequence and then carried out comparative analysis of LdPfn versus truncated LdPfn (δLdPfn) interactomes and functions in *L*. *donovani* promastigotes.

### Deletion of the **α**-helical insert from the LdPfn structure markedly affected its intracellular ligand binding affinity

As profilins are multi-ligand binding proteins [2], we carried out a comparative analysis of LdPfn versus δLdPfn interactomes. For this, we expressed GST conjugates of LdPfn and δLdPfn in *E. coli,* and purified the proteins (S2 Fig), as reported earlier [3]. The ligand binding profiles of LdPfn and δLdPfn were mapped by affinity pull-down and mass spectrometry, as described earlier [7]. Proteins that were identified in the affinity pull-down eluates of LdPfn and δLdPfn by mass spectrometry have been listed in S3 Table. About 200 proteins could be detected in eluates of GST-LdPfn affinity pull-down. Amongst which about 190 proteins were detected in 2-3 eluates of GST-LdPfn affinity pull-down as opposed to only about 140 proteins could be identified in 2-3 GST-δLdPfn pull-down eluates. Further, about 20 proteins including actin, mitochondrial outer membrane protein porin and calpain-like cysteine peptidase, that were identified in GST-LdPfn affinity pull-down eluates were completely missing in all the three eluates of GST-δLdPfn affinity pull -down (Table 1). These results clearly demonstrate that the intracellular ligand binding ability of LdPfn is markedly affected by deleting the helical insert region in the LdPfn structure.

**Table 1.** *L. donovani* proteins identified in eluates of three affinity pull-down assays with GST-LdPfn but not found in eluates of three affinity pull-down assays with GST-δLdPfn or GST alone, using mass spectrometry.

| Accession No. | Description |
| --- | --- |
| LdBPK_323110.1 | Nucleoside diphosphate kinase b |
| LdBPK_332040.1 | Phosphoribosyl pyrophosphate |
| LdBPK_342240.1 | Ribosomal protein l35a, putative |
| LdBPK_211160.1 | Histone H2A |
| LdBPK_332470.1 | Succinyl-coA:3-ketoacid-coenzyme |
| LdBPK_281100.1 | Ribosomal protein S20, putative |
| LdBPK_140700.1 | Fatty acid elongase, putative |
| LdBPK_140910.1 | Calpain-like cysteine peptidase, putative |
| LdBPK_351820.1 | Hypothetical protein, conserved |
| LdBPK_201320.1 | Calpain-like cysteine peptidase, putative |
| LdBPK_091020.1 | Elongation factor-1 gamma |
| LdBPK_020430.1 | Mitochondrial outer membrane protein, **Porin**, putative |
| LdBPK_041250.1 | **Actin** |
| LdBPK_110350.1 | 14-3-3 protein 2, putative |
| LdBPK_320020.1 | nuclear segregation protein, putative |
| LdBPK_250940.1 | cyclophilin a, putative |
| LdBPK_353840.1 | 60S ribosomal protein L23, putative |
| LdBPK_180700.1 | citrate synthase, putative |

**Table 2.**
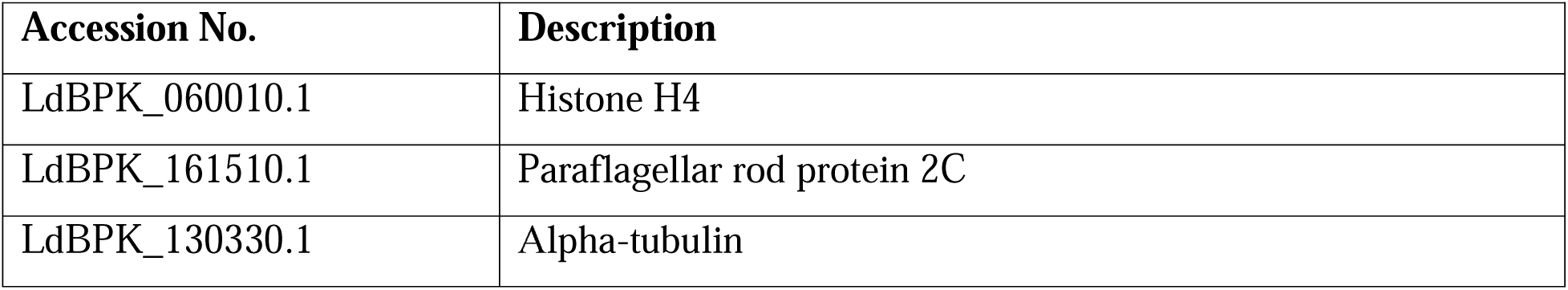
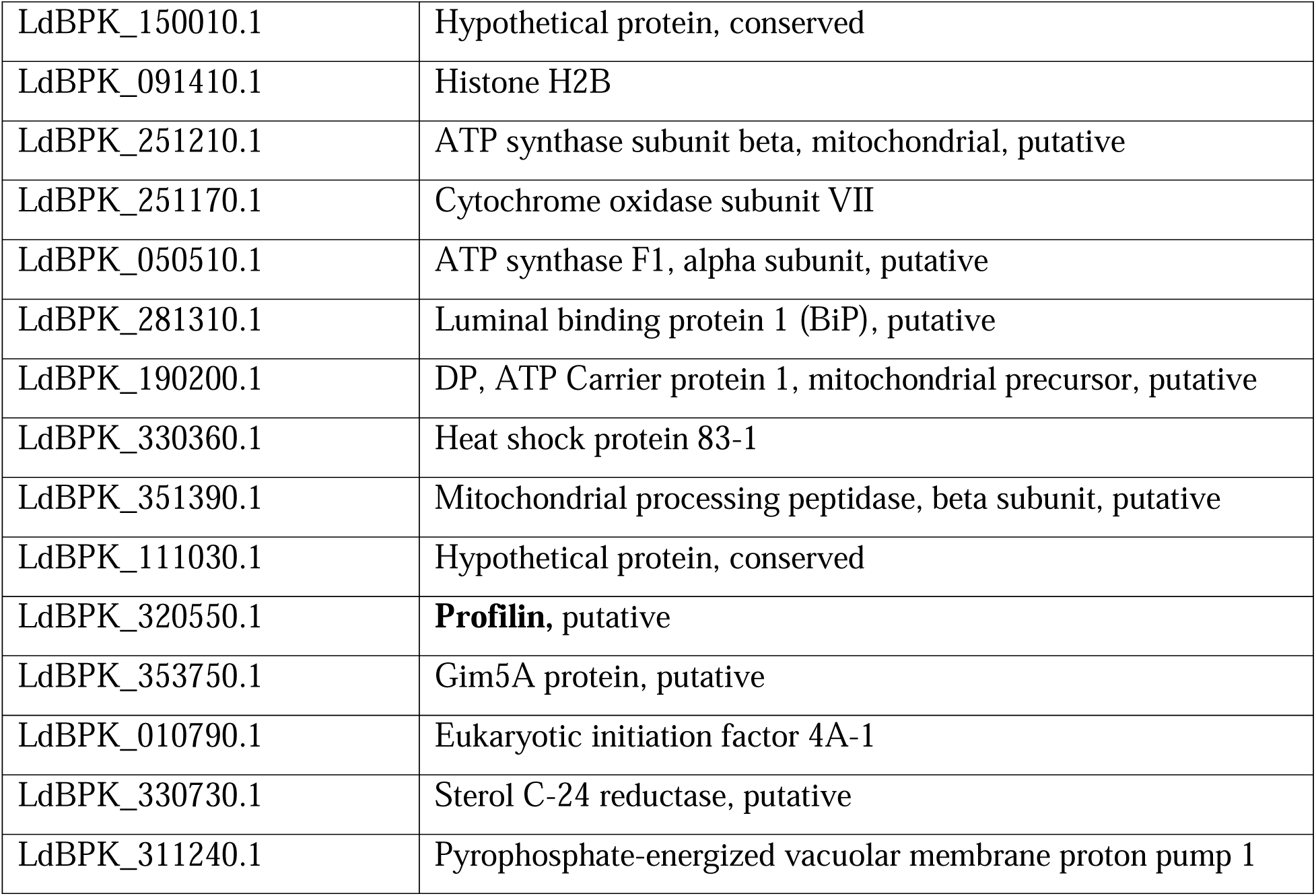
*L*. *donovani* proteins identified in eluates of two pull-down assays with GST-LdPorin but not found in eluates of two pull down assays with GST alone, using mass spectrometry.

As expected, deletion of α-helical insert region in the LdPfn structure should lead to loss of its ability to bind actin. This was confirmed by probing the pull-down eluates of both LdPfn and δLdPfn with anti-LdAct antibodies. Results given in Fig. 3 (A, B) clearly show that unlike LdPfn affinity pull-down eluates, anti-LdAct-antibodies failed to detect actin in eluates of the δLdPfn affinity pull-down.

**Fig. 3.**
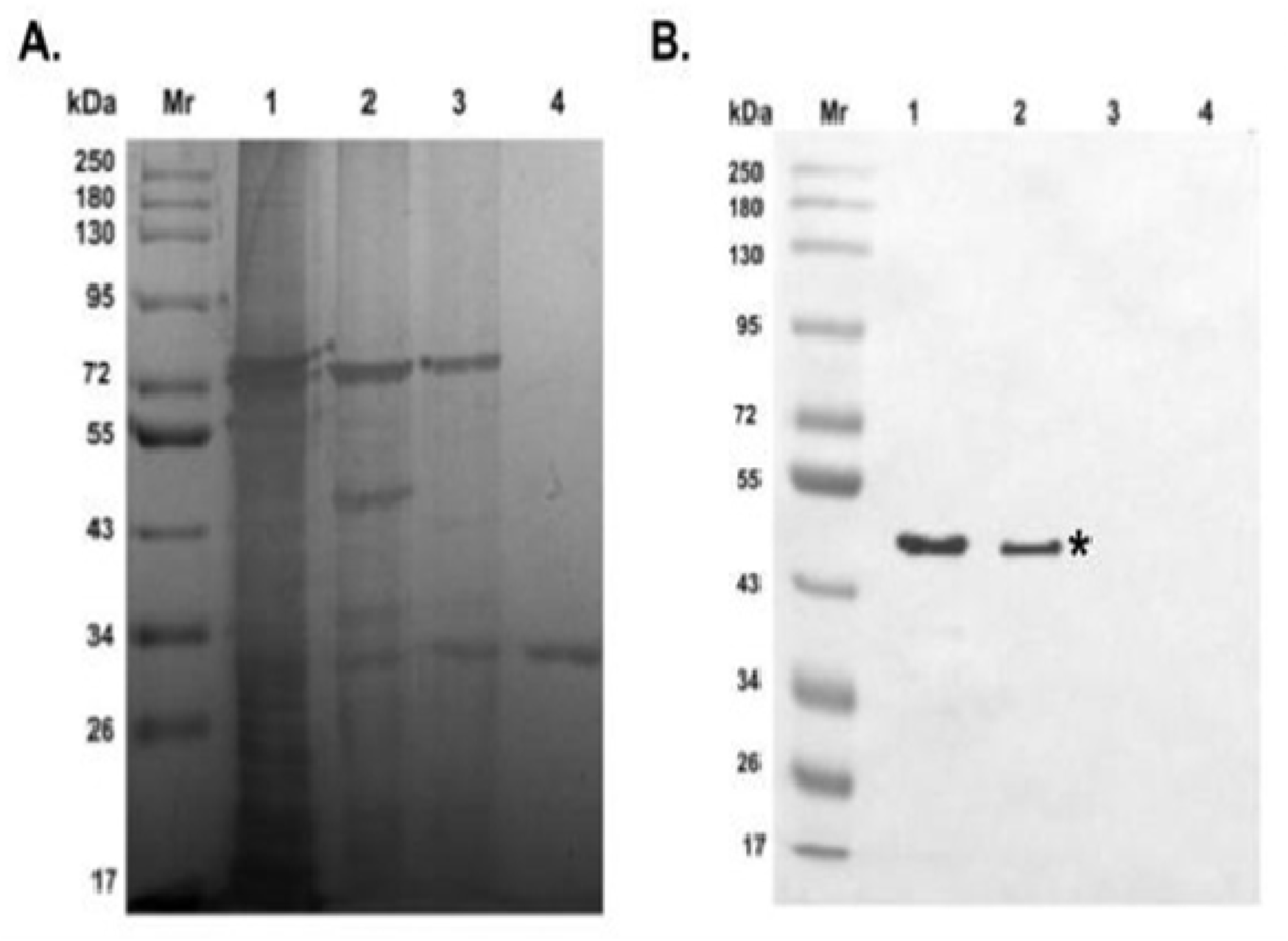
**(A) GST-LdPfn, GST-δ LdPfn and GST alone affinity pull-down eluates resolved on 12% polyacrylamide gel and stained with silver nitrate.** Mr, Molecular weight markers; Lane 1, *L. donovani* cell lysate; Lane 2, GST-LdPfn affinity pull-down eluate; Lane 3: GST-δLdPfn affinity pull-down eluate; Lane 4: GST alone affinity pull-down eluate. **(B) Western blot of ‘A’ using anti-LdAct antibodies**. Mr, Molecular weight markers; Lane 1, Total cell lysate; Lane 2, GST-LdPfn affinity pull-down eluate; Lane 3; GST-δLdPfn affinity pull down eluate; Lane 4: GST alone affinity pull-down eluate. Asterisk marks the position of LdAct.

As may be seen in Table 1, besides actin, the mitochondrial outer membrane protein, porin, was also missing in the GST-δLdPfn pull-down eluates. To further examine the validity of this finding, we cloned and purified the GST conjugate of porin protein (S3 Fig) and incubated it with Glutathione-Sepharose affinity beads at 37^0^ C for 2 h. The unbound protein was removed by centrifugation and after 2-3 washings, the bead-bound GST-Porin was incubated with clear lysates *of L. donovani* promastigotes at 4^0^ C overnight. After removing the unbound proteins by washing, the bead-bound proteins were eluted by boiling the beads in SDS-PAGE buffer at 96^0^C for 5 min. The eluted protein samples were resolved on SDS-polyacrylamide gel (12%) by electrophoresis. Two replicates of each of GST-Porin and GST alone pull-down samples were rinsed three times with water. The respective lanes were excised and submitted at the mass spectrometry facility for proteomic analysis. The proteomics data has been submitted at Proteome X-change consortium (Accession Id - PXD059335). In total, 18 proteins were identified in the eluates of GST-Porin pull down (Table2). While several of these proteins appeared to be the mitochondrial enzymes or their subunits, one of them was profilin, which again suggested that LdPfn directly interacts with porin.

This was further confirmed by probing the eluate of GST-Porin pull-down with anti-rabbit LdPfn antibodies (Fig.4). These results taken together strongly indicate that LdPfn interacts with mitochondrial outer membrane protein, porin, and that this interaction is abolished upon deleting the helical insert region in the LdPfn structure.

**Fig. 4.**
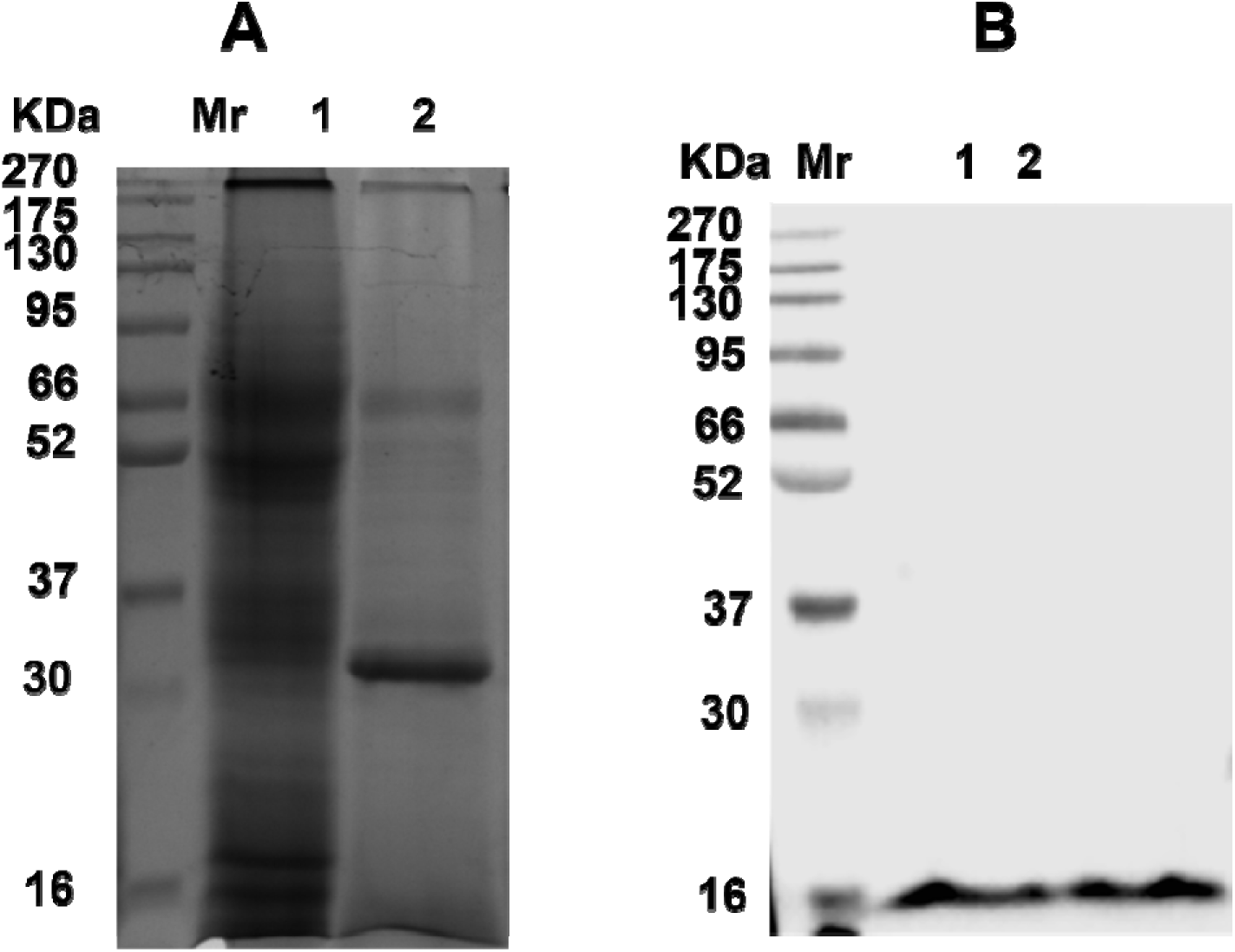
**(A) GST-LdPorin affinity pull-down eluate resolved on 12% polyacrylamide gel and stained with silver nitrate**. Mr, Molecular weight markers; Lane 1, *L. donovani* cell lysate; Lane 2, GST-LdPorin affinity pull-down eluate. **(B) Western blot of ‘A’ using anti-LdPfn antibodies**. Mr, Molecular weight markers; Lane 1, Total cell lysate; Lane 2, GST-LdPorin affinity pull-down eluate.

To examine whether the LdPfn-LdPorin interaction plays a role in regulation of the *Leishmania* mitochondrial functions, we measured cellular ATP levels in the wildtype cells expressing GFP-LdPfn and GFP-δLdPfn proteins. Fig.5 shows that ATP levels were significantly increased in the transgenic cells, as compared to wild-type control cells, suggesting that profilin-mediated actin remodeling could be involved in regulation of *Leishmania* mitochondrial functions.

**Fig. 5.**
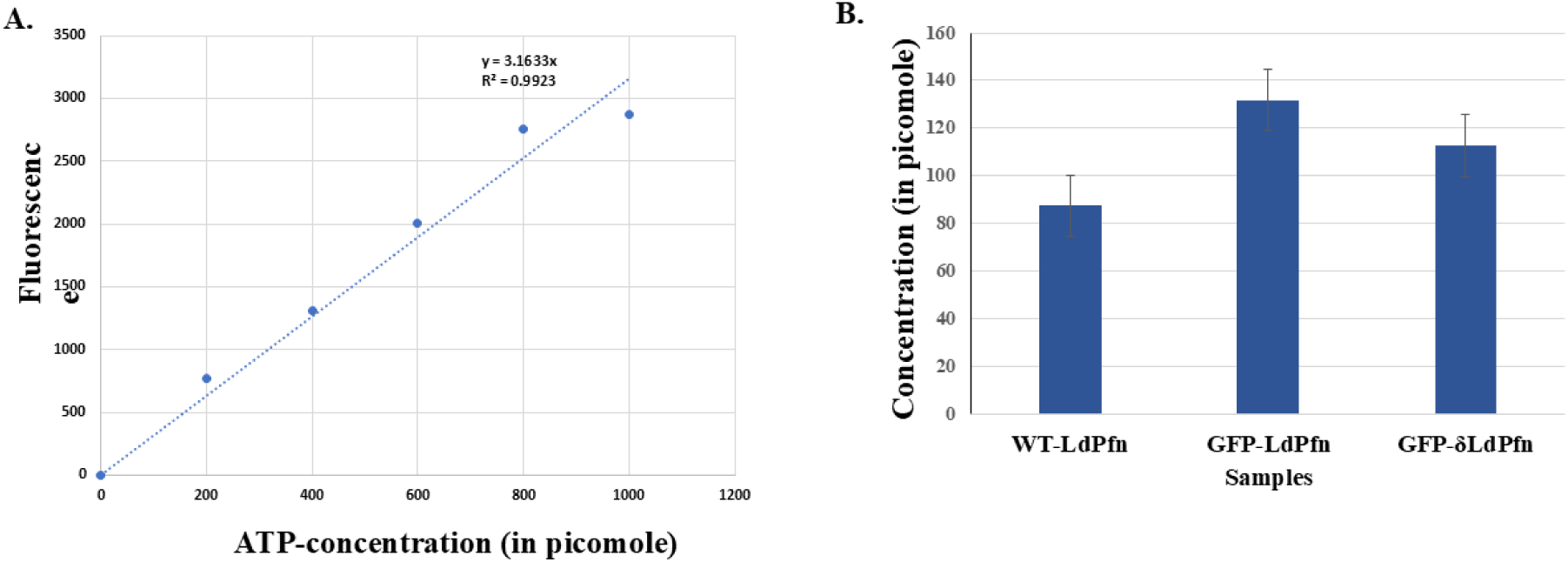
ATP levels in wild-type *Leishmania* cells were significantly increased by transfecting them with GFP-LdPfn or GFP-δLdPfn gene constructs. **(A)** ATP Standard curve **(B)** Cellular ATP levels. WT-LdPfn, wild-type cells; GFP-LdPfn, wild-type cells transfected with GFP-LdPfn gene construct; GFP-δLdPfn, wild-type cells transfected with GFP-δLdPfn gene construct. Results shown are means of three independent experiments ± SD. P-values: WT-LdPfn vs GFP-LdPfn, < 0.0005; WT-LdPfn vs GFP-δLdPfn, < 0.01.

### Deletion of the **α**-helical insert region in LdPfn structure leads to its reduced binding affinity to PLP motif containing proteins

Equal amounts of both GST-LdPfn and GST-δLdPfn were separately loaded on PLP-coupled Sepharose 4B columns. The columns were first washed with Tris -buffered 4M urea, and then the column – bound GST-LdPfn and GST-δLdPfn were eluted with Tris-buffered 8 M urea, as described in Material and Methods Section. Results given in Fig. 5 indicate that binding affinity of LdPfn to PLP motives is reduced by about 30% upon deleting the α-helical insert region in the LdPfn structure.

### Deletion of the **α**-helical insert region in the LdPfn structure adversely affected its functions

As LdPfn is known to play an important role in intracellular vesicle trafficking [3] and cell division cycle [7] in *Leishmania* cells, we examined whether these functions of LdPfn are affected by deleting the α-helical insert region in the LdPfn structure. For this, we transfected the wild type *Leishmania* cells separately with GFP-LdPfn and GFP-δLdPfn gene constructs, and then analyzed the growth, intracellular vesicle trafficking activity and cell division cycle in the transfected cells, as compared to non-transfected wild type cells. Whereas the growth of wild type *Leishmania* cells (Mean generation time:26h) was significantly slowed down (Mean generation time:35h) by transfecting them with GFP-δLdPfn gene construct, there was only little effect on the growth of these cells (Mean generation time:27.4 h) by transfecting them with GFP-LdPfn gene construct (Fig.6). These results suggest that direct binding of LdPfn with actin might have been playing an important role in driving the growth to the optimum level.

**Fig. 6.**
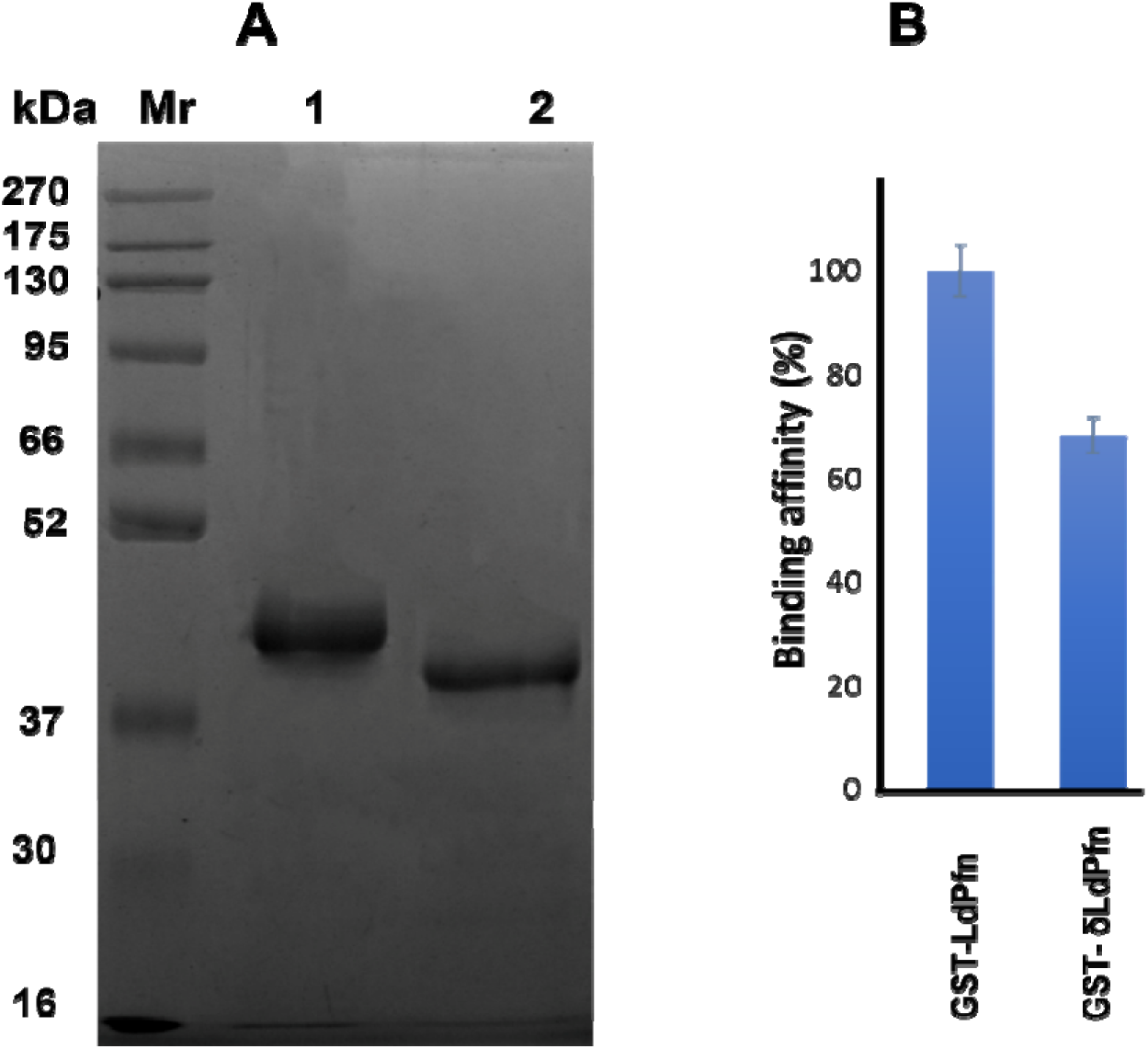
Deletion of the actin-binding domain in LdPfn amino acid sequence leads to its reduced binding to PLP motives. **(A) Binding of GST-LdPfn and GST-δLdPfn to poly-L-proline Sepharose**. Equal amount of purified GST-LdPfn and GST-δLdPfn proteins were subjected to affinity chromatography on poly-L-proline coupled to cyanogen bromide-activated Sepharose 4B beads. Mr, Molecular weight markers; lane 1, GST-LdPfn and lane 2, GST-δLdPfn. **(B) Quantitative analysis of poly-L-proline binding affinity of GST-LdPfn and GST-**δ**LdPfn.** Results shown are means of three independent experiments ± SD. P-value: GST-LdPfn vs GST-δLdPfn, < 0.00004.

### The intracellular vesicle trafficking activity of wild type Leishmania cells is decreased by transfecting them with GFP-**δ**LdPfn gene construct

To determine the effect of transfection of wild type *Leishmania* cells with GFP-LdPfn and GFP-δLdPfn gene constructs on their intracellular vesicle trafficking activity, we incubated the transfected cells first with a lipid soluble fluorescent dye FM4-64 FX [15], and then in fresh medium for varying time intervals. The incubated cells, after washing, were collected at various time points, spread on cover slips and then fixed with 2% paraformaldehyde. The fixed cells, after washing, were mounted and images were captured on immunofluorescence microscope. Whereas in the non-transfected cells, the FM4-64 FX dye traversed through the flagellar pocket, endosome and reached the lysosome/multivesicular tubule (MVT) by 120 min. However, in identical conditions, in the cells that were transfected with GFP-δLdPfn gene construct the dye could traverse only up to the nucleus (Fig.7). From these results, we infer that direct binding of LdPfn to actin is essentially required for driving intracellular vesicle trafficking in *Leishmania* cells.

**Fig. 7.**
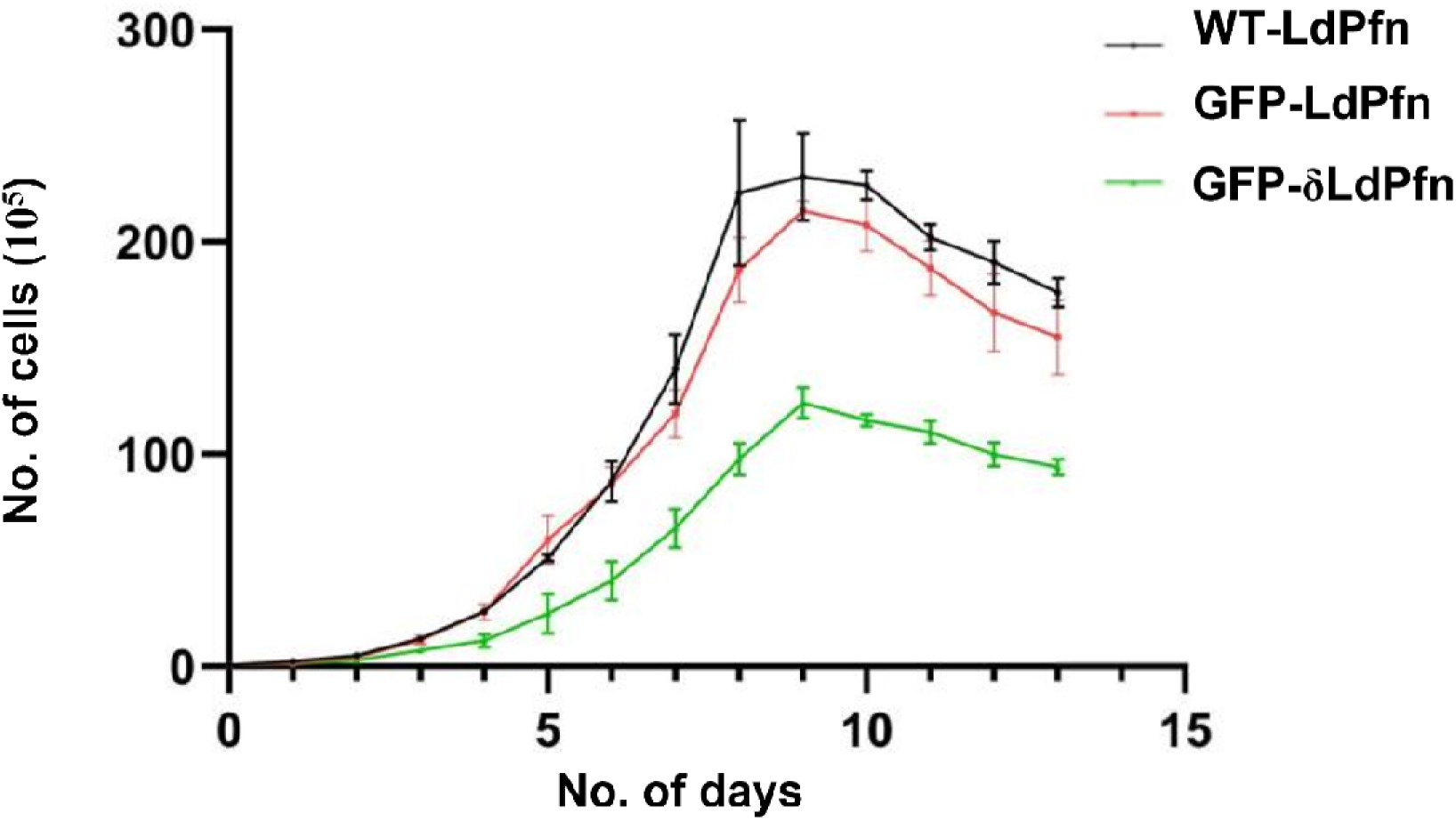
Growth of wild type *Leishmania* cells (black) is severely affected by transfecting them with GFP-δLdPfn (green) gene construct. The cells were grown in DMEM containing 10% FCS without antibiotics. Initial cell density was about 5 × 10^5^ cells ml^−1^. The growth wa recorded for 15 days. The values shown are means of three independent experiments± SD. Significant reduction (p-value < 0.02) in growth was seen in wild-type cells transfected with GFP-δLdPfn gene construct.

### Cell division cycle of wild type *Leishmania* cells is adversely affected by transfecting them with GFP-**δ**LdPfn gene construct

The effect of transfection of wild type *Leishmania* cells with GFP-LdPfn and GFP-δLdPfn gene constructs on their cell division cycle analyzed by employing flow cytometry. For this, the wild type *Leishmania* cells transfected with GFP-LdPfn and GFP-δLdPfn gene constructs were synchronized by treating them with N-hydroxyurea (HU) for 12 h. Approximately 70% of the cells were synchronized at the G1/S border. After release of the HU block, the cells were stained with propidium iodide (PI) to probe the total DNA content. The samples thus prepared were processed for the cell cycle analysis by flow cytometry. The wild type cells immediately entered from the G1- to -S phase, attaining its peak at 2 h and G2/M peak at 4h. However, both the G1- to-S and S-to-G2/M phase transitions in the wild type cells transfected with GFP-δLdPfn gene construct appeared to be delayed at least by 4h, as about 96% of these cells appeared at 6h in the S-phase and 60% cells at 8h in the G2/M phase (Fig.8A & B). Intriguingly, the wild type cells that were transfected with GFP-LdPfn gene construct, the G1-to-S phase and S-to-G2/M phase were delayed by 2h and 4h, respectively, compared to non -transfected wild type cells.

**Fig. 8.**
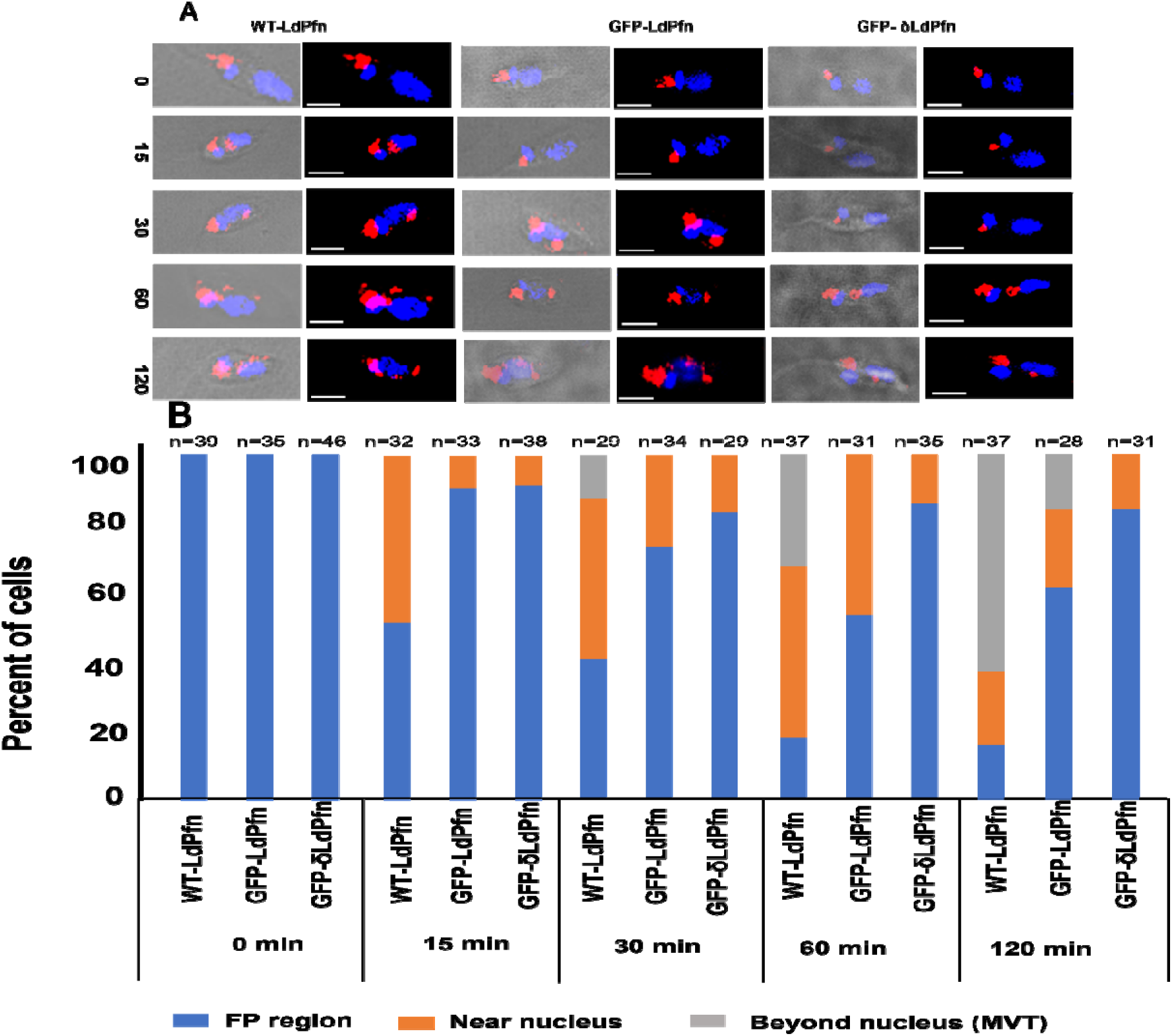
**(A) Intracellular trafficking of FM4-64 FX dye in wild type *Leishmania* cells is adversely affected by transfecting them with GFP-LdPfn and GFP-δLdPfn gene constructs.** The cells were first incubated in culture medium with 10% FCS and 2 μg/mL FM4-64 FX for 10 min at 25^0^C in the dark and then in fresh medium for varying time intervals. After washing with PBS, the cells collected at various time points were spread onto coverslips, fixed with 2% paraformaldehyde, washed, mounted and images were captured on immunofluorescenc microscope. In WT-LdPfn cells, the FM4-64 FX dye traversed through the flagellar pocket, endosome and reached the lysosome/multivesicular tubule (MVT) by 120 min, where as in GFP-δLdPfn cells the dye stacked near to the nucleus. Scale bar: 2 μm. **(B) Quantitative analysis of the intracellular movement of FM4-64 FX at different time points**. Three major categories of FM4-64 FX localization in the endocytic pathway were observed: Flagellar pocket region (FP), near the nucleus and beyond the nucleus. Percent fraction of ‘n’ number of cells where in the dye traversed to a given point at a given time points was plotted and the number of cells counted are indicated on the top of the respective bars. WT-LdPfn, wildtype cells; GFP-LdPfn, wild-type cells transfected with GFP-LdPfn gene construct; GFP-δLdPfn, wild-type cells transfected with GFP-δLdPfn gene construct. In about 60% of WT-LdPfn cells, the dye reached near to the posterior tip of the parasite in 120 min, while in about 80% GFP-δLdPfn cells the dye remained stuck near to the kinetoplast.

**Fig. 9.**
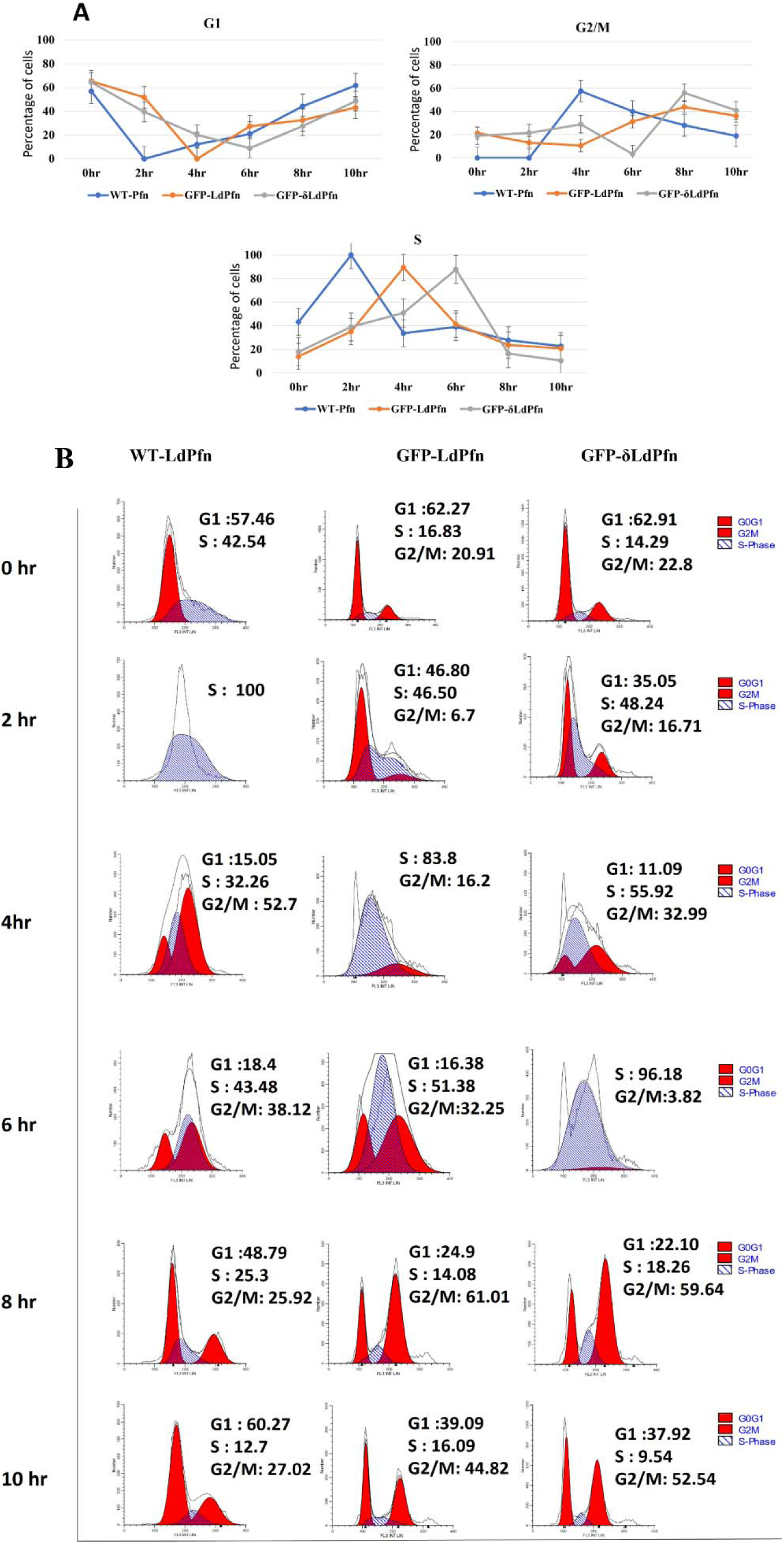
Cell division cycle of wild type *Leishmania* cells is altered after transfecting them with GFP-LdPfn and GFP-δLdPfn gene constructs. **(A)** Cell cycle distribution of wild type(WT-LdPfn) and wild type cells transfected with GFP-LdPfn and GFP-δLdPfn gene constructs, after removing the N-hydroxyurea block. Values shown are means of three experiments ± SD. **(B)** Representative flow cytometry data of wild type(WT-LdPfn) and wild type cells transfected with GFP-LdPfn and GFP-δLdPfn gene constructs.

## Discussion

Results presented here clearly show that deletion of the α-helical insert region (actin-binding domain) from the LdPfn amino acids sequence results in loss of its binding with more than 2 dozen proteins including actin, mitochondrial outer membrane protein, porin, and calpain-like cysteine peptidase. Also, the binding affinity of LdPfn to PLP motives is reduced by about 30%. It further revealed that expression of GFP-conjugate of δLdPfn in wild type *Leishmania* promastigotes leads to a marked reduction in their cell growth, decrease in intracellular vesicle trafficking activity and significantly delayed G1-to-S and S-to-G2/M phase transitions during their cell division cycle. As cellular processes such as growth, motility, intracellular trafficking, cell division, etc., require F-actin remodeling [17], wherein profilins play a key role [1,2, 18], we envisage that δLdPfn must have effectively competed out native LdPfn in its binding with actin along with other effector proteins/molecules, which in turn should have adversely affected actin remodeling-based intracellular activities in *Leishmania* cells.

F-Actin remodeling in *Leishmania* cells is primarily driven by profilin [3,8] together with actin depolymerizing factor, ADF/cofilin [19,20]. While LdPfn can regulate actin remodeling by virtue of its ability to promote or inhibit actin polymerization based on its intracellular concentration [3], ADF/cofilin binds and depolymerizes F-actin into ADP-bound G-actin, which after undergoing exchange of ADP for ATP by LdPfn, again recycled to the barbed end of growing actin filament [19, 20]. This apart, depletion of the intracellular pools of these two proteins have separately been shown to adversely affect the cell growth, intracellular vesicle trafficking and cell division cycle [3,7,19,21]. These studies taken together with the present study strongly indicate that F-actin remodeling plays an important role in regulation of growth, intracellular vesicle trafficking and cell division cycle in *Leishmania* promastigotes.

Besides actin, *Leishmania* profilin also binds to PLP stretches in proteins and polyphosphoinositides, such as PI (3,5) P2, PI (4,5) P2 and PI (3,4,5) P3 [3]. While PI (3,4,5) P3 and PI (4,5) P2 are generally found on the plasma membrane and their binding to profilin helps to regulate intracellular concentration of monomeric actin [22,23], PI (3,5) P2 is found predominantly within early and late endosomes or lysosomes and has been implicated to be involved in intracellular trafficking pathways [24]. Further, similarly to canonical profilins [25,26], *Leishmania major* profilin has recently been reported to recognize PLP stretches in proteins through its N- and C-terminal helices, distinctly away from the actin binding site [8]. This property of profilins has earlier been shown to enable them to bind vastly different partner proteins in multiple organisms [25, 26]. While several of these binding partners are involved in actin dynamics regulation, others are involved in regulation of endocytosis, nuclear export, Rac/Rho effector protein signaling [18] and possibly in some non-actin related pathways. As binding affinity of LdPfn to PLP stretches is largely retained even after deleting its actin-binding domain, we infer that besides direct binding of LdPfn to actin, its binding to other ligands, such as PI (3,5) P2 and PLP stretches in effector proteins also play an important role in regulation of intracellular functions of *Leishmania* profilins

Further, our earlier studies have suggested that LdPfn could play an important role in regulation of *Leishmania* mitochondrial functions through its interactions with the mitochondrial outer membrane protein, Porin (LdPorin) [7]. To further analyze this problem, we carried out comparative analysis of LdPfn and δLdPfn interactomes to find out whether LdPfn interacts with LdPorin independent of actin. Results revealed that interactions of LdPfn with LdPorin are abolished upon deleting the actin-binding domain in the LdPfn amino acid sequence, suggesting that LdPorin may be associated with dynamic actin filaments through LdPfn, which perhaps could be necessary for regulating mitochondrial positioning and functions. Mitochondria are doubly-membraned organelles that are fundamental to the reactive oxygen species (ROS) production, calcium homeostasis, cell proliferation or apoptosis [27], and are the main site to important metabolic reactions and ATP synthesis through the oxidative phosphorylation system [28]. The mitochondrial outer membrane contains several channel-forming proteins [29,30] including the major metabolite transporter protein, porin, that serves as an outer membrane channel for metabolites and as coupling factor for protein translocation into the inner membrane [30–33]. Interactions between the mitochondria and the cytoskeleton are essential for normal mitochondrial morphology, motility and distribution [34--38]. Emerging evidence indicates that besides microtubules and their motors, mitochondria also interact with the actin cytoskeleton in many cell types [34]. The actin cytoskeleton has been reported to play a multifaceted role in mitochondrial functions [34–39]. It not only influences the mitochondrial morphology, positioning, transport, metabolism and stress response [35–38], but it also regulates interactions of mitochondria with other cellular structures [35,39]. Proper mitochondrial positioning is crucial for efficient energy metabolism. Actin ensures that mitochondria are in the close proximity to areas of high energy demand [36], such as the flagellum and flagellar pocket in *Leishmania*. *Leishmania* cells contain only a single large mitochondrion, which is distributed in branches under the subpellicular microtubules, and in a specialized region, called kinetoplast, it houses its unusual genome, called kDNA [40]. The kinetoplast lies immediately adjacent to the flagellar basal bodies, and when the basal bodies replicate prior to cell division, the kinetoplast also replicates and segregates one mitochondrion to each daughter cell [41], suggesting that mitochondrion is perhaps tightly attached to the basal bodies. As actin and profilin are abundantly present in and around the kinetoplast including the basal bodies area and the flagellum [3, 42,43], we infer that the actin filaments may be serving as a linkage between the basal bodies and the kinetoplast. Besides this, actin filaments can interact with the mitochondrial outer membrane through the LdPfn binding to LdPorin, which would influence mitochondrial membrane potential and functions. That this indeed is the case, is well supported by our finding that the mitochondrial ATP levels were increased by expressing δLdPfn in wild-type *Leishmania* cells.

Finally, as revealed by the comparative interactome analysis, LdPfn appears to interact also with calpain-like cysteine proteases in an actin-dependent manner, suggesting that actin remodeling could be directly or indirectly involved in regulation of activity of these enzymes. Calpain-like cysteine peptidases are calcium-dependent multifunctional enzymes in *Leishmania*, which play a crucial role in parasite’s survival, virulence, and immune evasion [44]. These proteases contribute to the parasite’s ability to infect and survive within host macrophages, and have been implicated in the virulence of *Leishmania* parasites [45, 47, 49]. Also, they modulate host immune responses by cleaving host proteins involved in immune signaling, thereby aiding immune evasion [46, 48]. Further, calpain-like proteases being calcium-dependent, could act as key mediators of actin cytoskeletal changes in response to calcium fluctuations, which in turn would influence processes like cell motility and division [50,51]. We therefore speculate actin remodeling directly or indirectly may be involved in regulation of activities of these enzymes.

Lastly, the intracellular vesicle trafficking, cell division cycle, and cellular ATP levels were affected in the wild type *Leishmania* cells by transfecting them even with GFP-LdPfn. This means that these cellular activities of LdPfn are highly sensitive to its intracellular concentration, because both its depletion [3,7] or overexpression similarly affected these activities in *Leishmania* cells.

## Supporting information

All Supplementary data

## Acknowledgement

We are grateful to the Director, IBAB for providing us all possible support to conduct the present study, and thank the Centre for Cellular and Molecular Platforms (C-CAMP), Bangalore for Mass Spectrometric analysis.

## Data Availabilty

Mass spectrometry data related to LdPfn/ δLdPfn interactomes analysis have been deposited at PRIDE– Proteome X-change Consortium and may be accessed with the identifier: PXD039942. The mass spectrometry proteomics data of LdPorin have been deposited to the ProteomeXchange Consortium via the PRIDE partner repository with the dataset identifier PXD059335"

## Author Contributions

**Conceptualization**: Chhitar M. Gupta.

**Data Generation& Validation**: Sri kalaivani Raja &Prashant K. Rai

**Data Interpretation**: Sri kalaivani Raja, Prashant K. Rai & ChhitarM. Gupta

**Bioinformatics Analysis**: Hemant Kumar S. & S. Thiyagarajan

**Project Administration & Supervision**: Chhitar M. Gupta

**Writing**: Srikalaivani Raja & Chhitar M. Gupta

**Review & Editing**: Chhitar M. Gupta

## Supporting information

**S1_raw_images: The original, uncropped and unadjusted images of gels and their western blots.**

**S2Fig: (A) Coomassie blue stained 12% SDS-polyacrylamide gel showing purity of recombinant GST-LdPfn and GST-**δ**LdPfn.**Mr, Molecular weight markers; Lane 1, GST-LdPfn; Lane 2, GST-δLdPfn,.**(B)Western blots of purified LdPfn and** δ**LdPfn, employing anti-LdPfn antibodies.** Mr, Molecular weight markers; Lane 1, GST-LdPfn; Lane 2, GST-δLdPfn. Asterisk and arrow head mark the positions of GST-LdPfn and GST-δLdPfn, respectively

**S3Fig: (A) Coomassie blue stained 12% SDS-polyacrylamide gel showing purity of recombinant GST-LdPorin.** Mr, Molecular weight markers; Lane 1, GST-LdPorin. **(B) Western blot analysis of purified LdPorin, employing anti-GST antibodies.** Mr, Molecular weight markers; Lane 1, GST-LdPorin.

**S1Table: List of primers used for cloning** δ**LdPfn gene with GST tag at its N-terminus.**

**S2Table: List of primers used in cloning of mitochondrial outer membrane protein, porin, gene with GST tag at its N-terminus.**

**S3Table: List of proteins identified in 1-3 eluates of affinity pull-down with GST-LdPfn and GST-**δ**LdPfn by mass spectrometry**

