## Supplementary material for "Loss of direct binding of *Leishmania* profilin with actin adversely affected its functions and interactions with other cellular proteins": All Supplementary data

**Supporting information**

**S1Table: List of primers used for cloning δLdPfn gene with GST tag at its N-terminus.**

| **Construct** | **Primers** |
| --- | --- |
| δLdPfn-Forward | **5’GTGAACCCTGCGGACTGCAACAC 3’** |
| δLdPfn-Reverse | **5’GGGGTTCCCATATACGGCAACGATG 3’** |

**S2Table: List of primers used in cloning of mitochondrial outer membrane protein, porin, gene with GST tag at its N-terminus.**

| **Primer** | **Sequences** |
| --- | --- |
| Forward Primer | ATAGGATCCATGCCGCATTTGCAATCCGCC |
| Reverse Primer | TTATCTCGAGCTAAGAGTGCGTCATCTGCACG |

**S3Table: List of proteins identified in 1-3 eluates of affinity pull-down with GST-LdPfn and GST-δLdPfn by mass spectrometry**

| **Accession** | **Mass** | **Description** | GST-LdPfn | GST-δLdPfn |
| --- | --- | --- | --- | --- |
| LdBPK_170170.1 | 49097 | elongation factor 1-alpha | 3 | 3 |
| LdBPK_130330.1 | 49727 | alpha tubulin | 3 | 3 |
| LdBPK_283000.1 | 71224 | heat-shock protein hsp70, | 3 | 3 |
| LdBPK_281310.1 | 71786 | luminal binding protein 1 (BiP), | 2 | 2 |
| LdBPK_330360.1 | 80499 | heat shock protein 83-1 | 3 | 3 |
| LdBPK_081290.1 | 15882 | beta tubulin (fragment) | 3 | 3 |
| LdBPK_353150.1 | 99842 | ATP-dependent RNA helicase, | 3 | 3 |
| LdBPK_360210.1 | 94087 | elongation factor 2 | 3 | 3 |
| LdBPK_010790.1 | 45299 | Eukaryotic initiation factor 4A-1 | 3 | 3 |
| LdBPK_362140.1 | 59305 | chaperonin HSP60, mitochondrial precursor | 3 | 3 |
| LdBPK_362130.1 | 60517 | chaperonin HSP60, mitochondrial precursor | 3 | 2 |
|  | 46007 | Enolase \| | 3 | 3 |
| LdBPK_230410.1 | 38433 | NADP-dependent alcohol | 3 | 3 |
| LdBPK_040750.1 | 24541 | 60S ribosomal protein L10, | 3 | 3 |
| LdBPK_141240.1 | 115050 | glutamate dehydrogenase \| | 3 | 3 |
| LdBPK_311930.1 | 14673 | ubiquitin-fusion protein \| | 3 | 3 |
| LdBPK_340870.1 | 25920 | elongation factor 1-beta | 2 | 1 |
| LdBPK_320410.1 | 66767 | ATP-dependent RNA helicase | 3 | 3 |
| LdBPK_312890.1 | 20162 | ADP-ribosylation factor, putative | 3 | 1 |
| LdBPK_343440.1 | 18008 | 60S ribosomal protein L21, | 3 | 3 |
| LdBPK_201350.1 | 15016 | small myristoylated protein 1 | 3 | 3 |
| LdBPK_282970.1 | 34351 | activated protein kinase c receptor | 3 | 2 |
| LdBPK_363020.1 | 15760 | 40S ribosomal protein S24e | 3 | 2 |
| LdBPK_320550.1 | 16135 | profilin, putative | 3 | 3 |
| LdBPK_091410.1 | 12264 | histone H2B | 3 | 3 |
| LdBPK_332520.1 | 71968 | heat shock protein, putative | 3 | 2 |
| LdBPK_060030.1 | 75400 | WD domain, G-beta repeat, | 3 | 2 |
| LdBPK_070550.1 | 39077 | 60S ribosomal protein L7a, | 3 | 3 |
| LdBPK_365353.1 | 27510 | 40S ribosomal protein SA, | 3 | 2 |
| LdBPK_323110.1 | 16613 | nucleoside diphosphate kinase b | 3 | 0 |
| LdBPK_050510.1 | 62509 | ATP synthase F1, alpha subunit, putative | 3 | 3 |
| LdBPK_330960.1 | 37717 | 40S ribosomal protein S3, | 3 | 3 |
| LdBPK_302480.1 | 54172 | heat shock 70-related protein 1, mitochondrial | 3 | 3 |
| LdBPK_261220.1 | 70394 | heat shock protein 70-related protein | 3 | 2 |
| LdBPK_354200.1 | 64604 | polyadenylate-binding protein | 3 | 2 |
| LdBPK_161510.1 | 68859 | paraflagellar rod protein 2C | 3 | 3 |
| LdBPK_130460.1 | 15582 | 40S ribosomal protein S12, | 3 | 3 |
| LdBPK_260840.1 | 16677 | 40S ribosomal protein S16, | 3 | 3 |
| LdBPK_361310.1 | 22124 | 40S ribosomal protein S9 | 3 | 3 |
| LdBPK_261960.1 | 88994 | hypothetical protein, conserved | 3 | 3 |
| LdBPK_331690.1 | 36228 | 3-ketoacyl-CoA reductase, | 3 | 3 |
| LdBPK_290790.1 | 86902 | heat shock protein 90 | 3 | 1 |
| LdBPK_251210.1 | 56257 | ATP synthase subunit beta, | 3 | 3 |
| LdBPK_332040.1 | 39010 | phosphoribosylpyrophosphate | 3 | 0 |
| LdBPK_353280.1 | 44498 | cystathione gamma lyase, putative | 3 | 2 |
| LdBPK_350600.1 | 20776 | 60S ribosomal protein L18a, | 3 | 3 |
| LdBPK_081020.1 | 62200 | stress-induced protein sti1 | 2 | 3 |
| LdBPK_350400.1 | 29989 | 40S ribosomal protein S3A, | 3 | 3 |
| LdBPK_291890.1 | 69016 | paraflagellar rod protein 1D, | 3 | 2 |
| LdBPK_367320.1 | 81810 |  | 3 | 2 |
| LdBPK_111180.1 | 14687 |  | 3 | 3 |
| LdBPK_261950.1 | 286879 | microtubule-associated protein, | 3 | 2 |
| LdBPK_342240.1 | 16444 | ribosomal protein l35a, putative | 3 | 0 |
| LdBPK_190200.1 | 35131 | ADP, ATP carrier protein 1, | 3 | 3 |
| LdBPK_070130.1 | 59588 | ATP-dependent RNA helicase | 3 | 1 |
| LdBPK_260160.1 | 28841 | 60S ribosomal protein L7, | 3 | 3 |
| LdBPK_282480.1 | 46391 | eukaryotic translation initiation | 3 | 1 |
| LdBPK_230860.1 | 47321 | 3-ketoacyl-CoA thiolase-like | 3 | 3 |
| LdBPK_111110.1 | 16361 | 60S ribosomal protein L28, | 1 | 2 |
| LdBPK_230580.1 | 78121 | acetyl-CoA synthetase, putative | 2 | 2 |
| LdBPK_211160.1  LdBPK_292620.1 | 13927 | histone H2A | 3 | 0 |
| LdBPK_364730.1 | 22008 | 60S ribosomal protein L18, | 3 | 3 |
| LdBPK_131120.1 | 30631 | 40S ribosomal protein S4, | 3 | 2 |
| LdBPK_332470.1 | 52596 | succinyl-coA:3-ketoacid-coenzyme | 3 | 0 |
| LdBPK_150270.1 | 67146 | lysyl-tRNA synthetase, putative | 3 | 2 |
| LdBPK_151120.1 | 12672 | tryparedoxin peroxidase | 2 | 2 |
| LdBPK_230050.1 | 25368 | peroxidoxin | 3 | 2 |
| LdBPK_311130.1 | 43984 | amidohydrolase, putative | 2 | 1 |
| LdBPK_363950.1 | 8893 | 60S ribosomal protein L10a, | 3 | 2 |
| LdBPK_312210.1 | 31607 | prostaglandin f2-alpha synthase \| | 3 | 3 |
| LdBPK_211290.1 | 21516 | 60S ribosomal protein L9, | 3 | 3 |
| LdBPK_290860.1 | 36995 | cysteine peptidase C (CPC) | 3 | 2 |
| LdBPK_323200.1 | 96090 | hypothetical protein, conserved | 3 | 2 |
| LdBPK_251220.1 | 13038 | ribosomal protein S25 | 3 | 1 |
| LdBPK_272350.1 | 43620 | heat shock protein DNAJ, putative | 3 | 2 |
| LdBPK_170010.1 | 87550 | eukaryotic translation initiation | 3 | 2 |
| LdBPK_281100.1 | 13003 | ribosomal protein S20, putative | 3 | 0 |
| LdBPK_180740.1 | 51490 | Elongation factor Tu, | 3 | 2 |
| LdBPK_260780.1 | 19404 | glutathione peroxidase-like | 2 | 0 |
| LdBPK_302570.1 | 22097 | reticulon domain protein, 22 kDa | 2 | 1 |
| LdBPK_352240.1 | 30239 | RNA-binding protein, putative | 3 | 1 |
| LdBPK_351240.1 | 28100 | short chain dehydrogenase | 1 | 0 |
| LdBPK_364100.1 | 47737 | S-adenosylhomocysteine | 3 | 2 |
| LdBPK_252100.1 | 33049 | hypothetical protein, conserved | 2 | 1 |
| LdBPK_220340.1 | 7461 | 40S ribosomal protein S15, | 3 | 3 |
| LdBPK_100210.1 | 52579 | nucleolar protein 56, putative | 2 | 1 |
| LdBPK_320790.1 | 25194 | nuclear RNA binding domain | 3 | 3 |
| LdBPK_260610.1 | 10667 | 10 kDa heat shock protein, | 2 | 2 |
| LdBPK_100960.1 | 23388 | small GTP-binding protein Rab11, | 1 | 1 |
| LdBPK_270620.1 | 22256 | ras-related protein RAB1A, | 2 | 2 |
| LdBPK_181350.1 | 91683 | heat shock protein, putative \| | 3 | 2 |
| LdBPK_241700.1 | 66698 | Succinate dehydrogenase | 2 | 0 |
| LdBPK_362790.1 | 48605 | dihydrolipoamide | 3 | 2 |
| LdBPK_100310.1 | 48463 | isocitrate dehydrogenase [NADP], mitochondrial precurs | 2 | 2 |
| LdBPK_060010.1 | 11417 | histone H4 | 3 | 2 |
| LdBPK_303560.1 | 43057 | S-adenosylmethionine synthetase | 3 | 2 |
| LdBPK_361700.1 | 190952 | clathrin heavy chain, putative | 2 | 2 |
| LdBPK_061320.1 | 71121 | pteridine transporter, putative | 2 | 2 |
| LdBPK_212190.1 | 10306 | 60S ribosomal protein L37a, | 2 | 0 |
| LdBPK_140700.1 | 32825 | fatty acid elongase, putative | 3 | 0 |
| LdBPK_281020.1 | 45540 | Mitochondrial import receptor | 2 | 1 |
| LdBPK_180510.1 | 97392 | aconitase, putative | 2 | 0 |
| LdBPK_281050.1 | 15583 | 40S ribosomal protein S14 | 3 | 3 |
| LdBPK_252520.1 | 109274 | hypothetical protein, conserved | 2 | 1 |
| LdBPK_271140.1 | 35282 | Serine-threonine kinase receptor-associated protein | 2 | 1 |
| LdBPK_160760.1 | 36930 | transaldolase, putative | 3 | 1 |
| LdBPK_100900.1 | 13878 | nuclear transport factor 2, | 3 | 1 |
| LdBPK_342530.1 | 28833 | eukaryotic translation initiation factor 3 subunit g | 3 | 1 |
| LdBPK_120490.1 | 67025 | glucose-6-phosphate isomerase | 1 | 0 |
| LdBPK_241560.1 | 19435 | IgE-dependent histamine-releasing factor, putative | 2 | 2 |
| LdBPK_351030.1 | 39755 | casein kinase, putative | 2 | 2 |
| LdBPK_291360.1 | 96897 | ATP-dependent Clp protease subunit, heat shock protein 100 (HSP100) | 2 | 1 |
| LdBPK_020680.1 | 90768 | ATP-dependent Clp protease subunit, heat shock protein 78 (HSP78), | 3 | 2 |
| LdBPK_321430.1 | 85808 | G5-interacting protein, putative | 3 | 1 |
| LdBPK_351390.1 | 54553 | mitochondrial processing peptidase, beta subunit, putative | 3 | 2 |
| LdBPK_041170.1 | 38832 | fructose-1,6-bisphosphatase, cytosolic, putative | 3 | 3 |
| LdBPK_060580.1 | 30249 | deoxyuridine triphosphatase, putative | 2 | 0 |
| LdBPK_060350.1 | 46029 | NAD(P)-dependent steroid dehydrogenase protein, putative | 2 | 0 |
| LdBPK_221370.1 | 25984 | 40S ribosomal protein L14, putative | 3 | 2 |
| LdBPK_301030.1 | 22940 | p22 protein precursor, putative | 2 | 0 |
| LdBPK_360990.1 | 18980 | transcript_product=40S ribosomal protein S18, putative \| | 2 | 2 |
| LdBPK_030190.1 | 61901 | delta-1-pyrroline-5-carboxylate dehydrogenase, putative | 3 | 2 |
| LdBPK_140910.1 | 12923 | calpain-like cysteine peptidase, putative | 3 | 0 |
| LdBPK_071150.1 | 38836 | RNA binding protein-like protein | 2 | 0 |
| LdBPK_221390.1 | 105914 | alanyl-tRNA synthetase, putative | 2 | 1 |
| LdBPK_352790.1 | 53474 | galactokinase-like protein | 2 | 1 |
| LdBPK_351820.1 | 81817 | hypothetical protein, conserved | 3 | 0 |
| LdBPK_322330.1 | 26288 | eukaryotic translation initiation factor 3 subunit k | 3 | 2 |
| LdBPK_340010.1 | 33475 | short chain dehydrogenase, putative | 2 | 2 |
| LdBPK_312320.1 | 38618 | 3,2-trans-enoyl-CoA isomerase, mitochondrial precursor, putative | 2 | 1 |
| LdBPK_171390.1 | 80760 | eukaryotic translation initiation factor 3 subunit b | 3 | 2 |
| LdBPK_201320.1 | 17025 | calpain-like cysteine peptidase, putative | 3 | 0 |
| LdBPK_150010.1 | 11417 | histone H4 | 3 | 1 |
| LdBPK_230310.1 | 30203 | pteridine reductase 1 | 2 | 1 |
| LdBPK_353900.1 | 61554 | T-complex protein 1, eta subunit, putative | 3 | 1 |
| LdBPK_111000.1 | 100883 | pyruvate phosphate dikinase, putative | 3 | 2 |
| LdBPK_091020.1 | 51279 | elongation factor-1 gamma | 3 | 0 |
| LdBPK_342410.1 | 22606 | ALBA-domain protein 3 | 1 | 1 |
| LdBPK_340570.1 | 65759 | Cysteine leucine rich protein | 1 | 0 |
| LdBPK_352380.1 | 99535 | Importin 1 | 1 | 1 |
| LdBPK_350110.1 | 20158 | coatomer subunit zeta, putative | 1 | 0 |
| LdBPK_364070.1 | 38502 | eukaryotic translation initiation factor 3 subunit, putative | 3 | 2 |
| LdBPK_300470.1. | 62507 | aspartyl-tRNA synthetase, putative | 1 | 2 |
| LdBPK_161500.1 | 39350 | hypothetical protein, conserved | 1 | 1 |
| LdBPK_141050.1 | 40131 | ADP/ATP mitochondrial carrier-like protein | 3 | 2 |
| LdBPK_161710.1 | 30211 | prohibitin | 2 | 2 |
| LdBPK_290950.1 | 20198 | ADP-ribosylation factor-like protein 3A, putative | 2 | 2 |
| LdBPK_261690.1 | 22268 | cytochrome c oxidase subunit V, putative | 3 | 3 |
| LdBPK_367280.1 | 52324 | protein disulfide isomerase 2 | 2 | 2 |
| LdBPK_072480.1 | 34757 | 60S acidic ribosomal protein P0, putative | 3 | 2 |
| LdBPK_230720.1 | 75198 | Kinesin-C | 2 | 2 |
| LdBPK_303480.1 | 90516 | protein mkt1, putative | 3 | 2 |
| LdBPK_361420.1 | 86764 | Valosin-containing protein, putative | 3 | 3 |
| LdBPK_353110.1 | 126366 | ubiquitin-activating enzyme E1, putative | 2 | 1 |
| LdBPK_282100.1 | 18674 | ribose 5-phosphate isomerase, putative | 1 | 1 |
| LdBPK_242320.1 | 19970 | hypothetical predicted multi-pass transmembrane protein | 2 | 2 |
| LdBPK_141460.1 | 74890 | tyrosyl-tRNA synthetase, putative | 3 | 2 |
| LdBPK_240040.1 | 19101 | 60S ribosomal protein L17, putative | 3 | 2 |
| LdBPK_282600.1 | 41748 | 2-oxoglutarate dehydrogenase, E2 component, dihydrolipoamide succinyltransferase, putative | 2 | 3 |
| LdBPK_365330.1 | 103915 | hypothetical protein, conserved | 3 | 2 |
| LdBPK_351190.1 | 122988 | NADH-dependent fumarate reductase, putative | 1 | 1 |
| LdBPK_020340.1 | 58966 | proteasome regulatory non-ATPase subunit 6, putative | 2 | 2 |
| **LdBPK_020430.1** | **31905** | **Mitochondrial outer membrane protein porin, putative** | **3** | **0** |
| LdBPK_140190.1 | 22392 | Thioredoxin-like, putative | 2 | 2 |
| LdBPK_290640.1 | 206739 | ATP-binding cassette protein subfamily A, member 10, putative | 1 | 1 |
| LdBPK_352250.1 | 11228 | kinetoplastid membrane protein-11 | 2 | 0 |
| LdBPK_292310.1 | 77968 | dynamin-1-like protein | 3 | 2 |
| LdBPK_231460.1 | 60163 | T-complex protein 1, gamma subunit, putative | 2 | 2 |
| LdBPK_180780.1 | 27151 | DNA-directed RNA polymerases II, putative | 1 | 1 |
| LdBPK_366760.1 | 41041 | mitogen activated protein kinase homologue | 3 | 2 |
| LdBPK_330660.1 | 14339 | paraflagellar rod component, | 1 | 0 |
| LdBPK_131320.1 | 16001 | ubiquitin-conjugating enzyme, putative | 2 | 0 |
| LdBPK_303130.1 | 35583 | RNA-binding protein 42 (RNA-binding motif protein 42), putative | 2 | 2 |
| LdBPK_071020.1 | 42182 | splicing factor ptsr1-like protein | 3 | 2 |
| LdBPK_321000.1 | 102024 | Staphylococcal nuclease homologue/Tudor domain containing protein, putative | 3 | 2 |
| LdBPK_200110.1 | 51295 | phosphoglycerate kinase C, glycosomal | 3 | 2 |
| LdBPK_120120.1 | 149600 | hypothetical protein, conserved | 3 | 1 |
| LdBPK_355300.1 | 39565 | isopentenyl-diphosphate delta-isomerase (type II), putative | 3 | 2 |
| LdBPK_281700.1 | 38280 | hydrolase, alpha/beta fold family, putative | 1 | 2 |
| LdBPK_360080.1 | 29022 | stress-inducible protein STI1 homolog | 3 | 2 |
| LdBPK_070710.1 | 37928 | eukaryotic translation initiation factor 3 subunit h | 3 | 1 |
| LdBPK_170870.1 | 51978 | GMP reductase | 1 | 0 |
| LdBPK_302860.1 | 11772 | succinate dehydrogenase subunit 3 | 1 | 0 |
| LdBPK_131260.1 | 90927 | hypothetical protein, conserved | 2 | 2 |
| LdBPK_220490.1 | 49471 | proteasome regulatory ATPase subunit 5, putative | 3 | 3 |
| LdBPK_352230.1 | 17513 | 60S ribosomal protein L12,putative | 3 | 2 |
| LdBPK_030670.1 | 246535 | DEAD/DEAH box helicase, putative | 3 | 3 |
| **LdBPK_041250.1** | **42022** | **actin** | **3** | **0** |
| LdBPK_170970.1 | 48353 | hypothetical protein, conserved | 3 | 2 |
| LdBPK_291190.1 | 40013 | Ankyrin repeats (3 copies), putative | 2 | 2 |
| LdBPK_181360.1 | 42692 | pyruvate dehydrogenase E1 component alpha subunit, putative | 2 | 2 |
| LdBPK_110350.1 | 29214 | 14-3-3 protein 2, putative | 3 | 2 |
| LdBPK_130270.1 | 13352 | ALBA-domain protein 1 | 1 | 1 |
| LdBPK_353030.1 | 51003 | chaperone protein DnaJ, putative | 2 | 2 |
| LdBPK_055450.1 | 55158 | pyruvate kinase, putative | 2 | 2 |
| LdBPK_271770.1 | 74197 | trypanothione synthetase | 3 | 2 |
| LdBPK_131400.1 | 58786 | chaperonin TCP20, putative | 3 | 1 |
| LdBPK_181510.1 | 107269 | P-type ATPase 1B, putative | 3 | 2 |
| LdBPK_030960.1 | 46610 | eukaryotic translation initiation factor 2 subunit alpha | 2 | 1 |
| LdBPK_212140.1 | 34373 | ATP synthase F1 subunit gamma protein, putative | 2 | 1 |
| LdBPK_342680.1 | 71409 | egulatory subunit of protein kinase a-like protein | 2 | 0 |

**S1Fig: The original, uncropped and unadjusted images of gels and their western blots.**
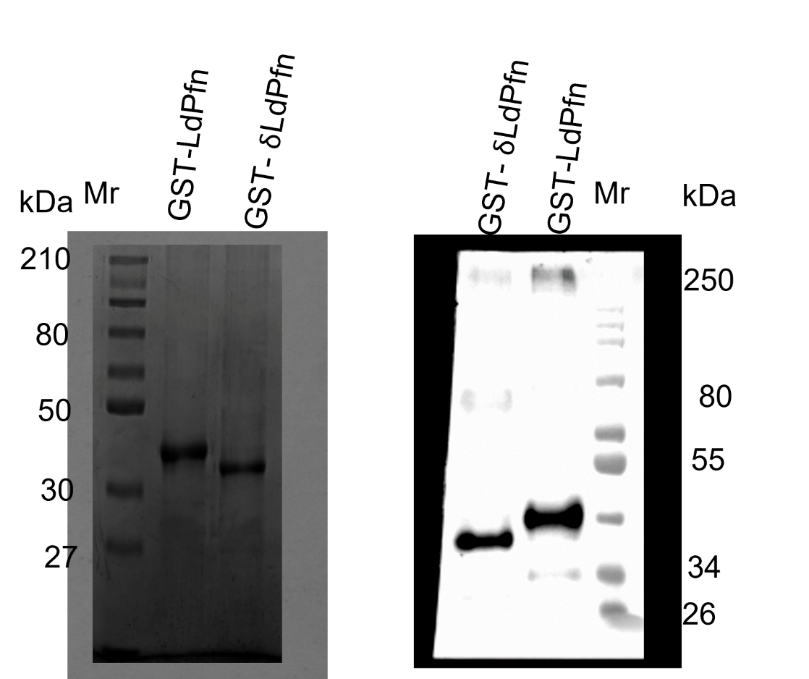
**Fig. 1**

Coomassie stained SDS PAGE Western blot with anti-GST antibody

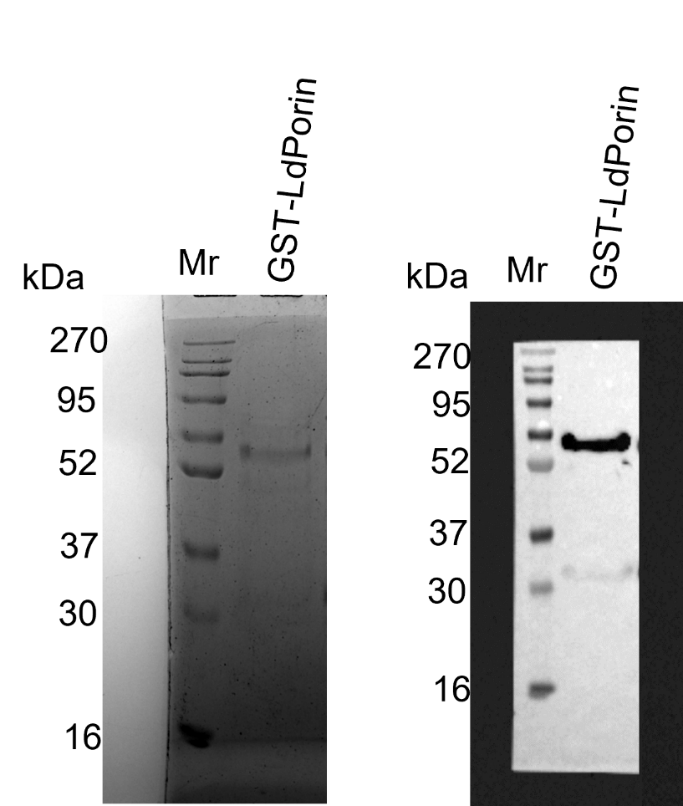
**Fig 2**

Coomassie stained SDS PAGE	Western blot with anti-GST antibodies

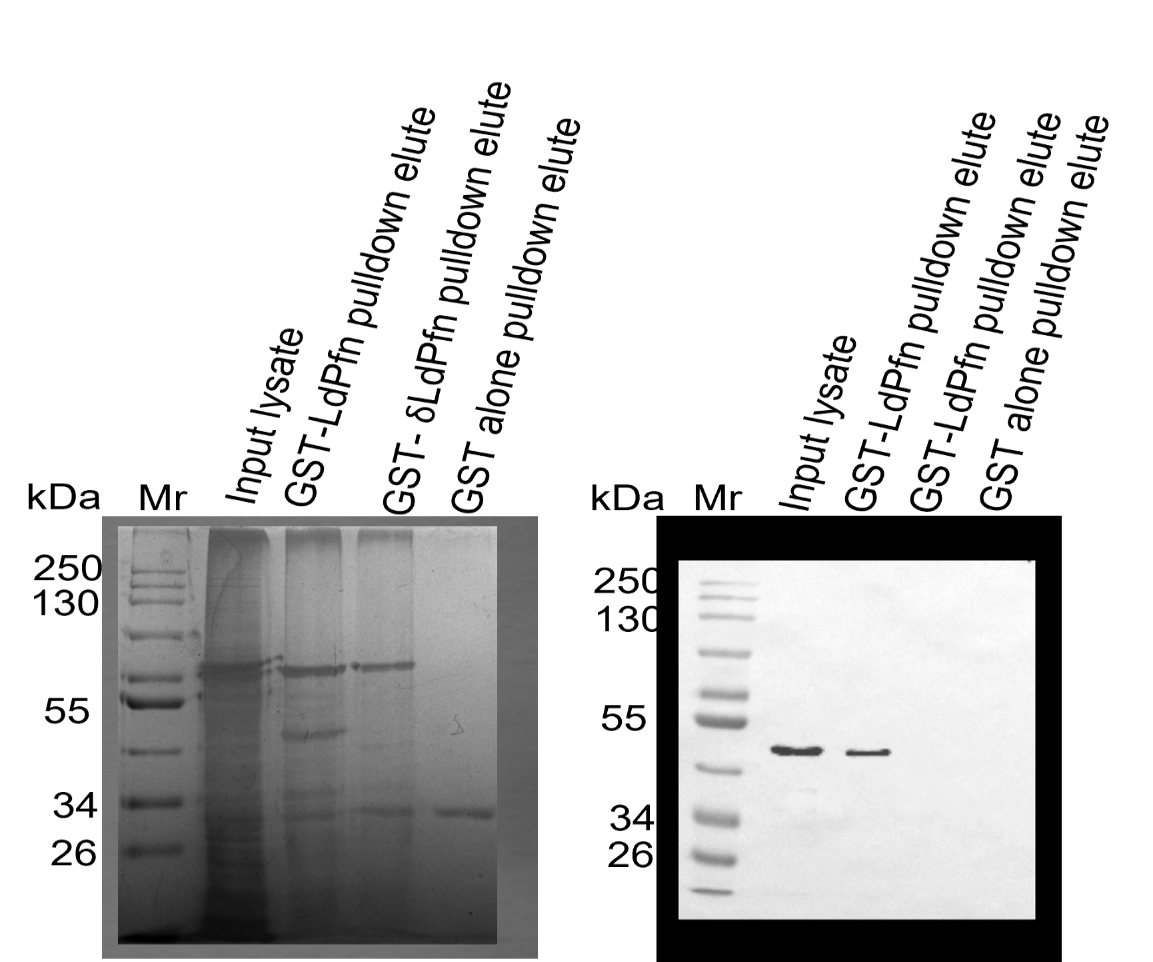
**Fig 3**

Silver-stained polyacrylamide gel	Western blot with anti-Ldactin antibodies

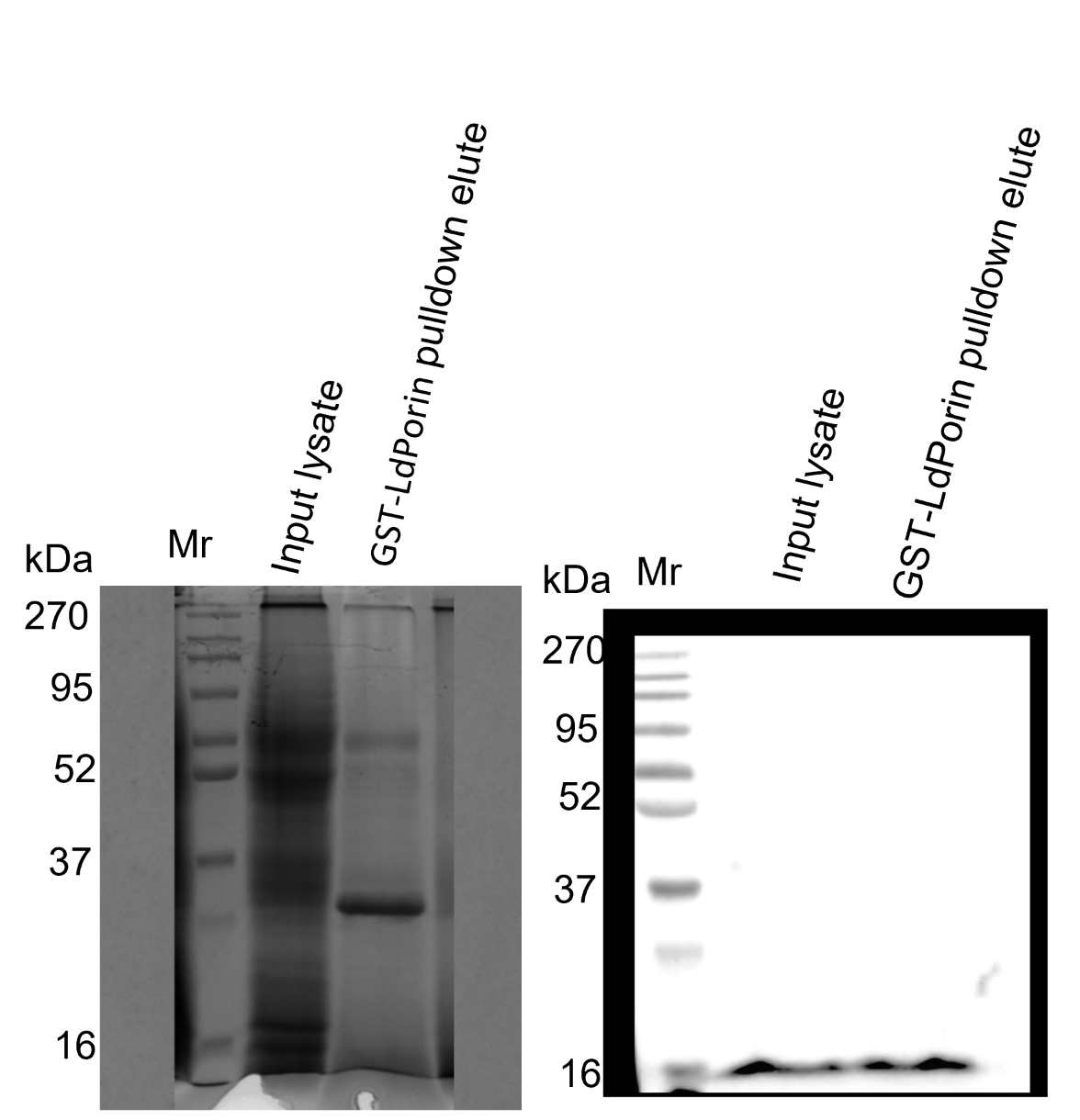
**Fig 4**

Silver-stained polyacrylamide gel	Western blot with anti-LdPfn antibodies

**Fig S3: The original, uncropped and unadjusted images of gels and their western blots.**

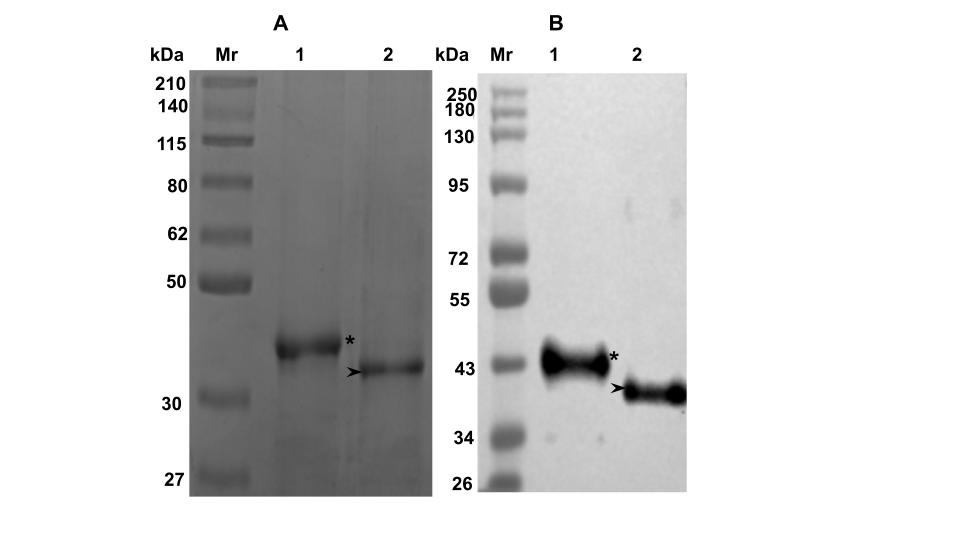

**S2Fig: (A) Coomassie blue stained 12% SDS-polyacrylamide gel showing purity of recombinant GST-LdPfn and GST-δLdPfn.**Mr, Molecular weight markers; Lane 1, GST-LdPfn; Lane 2, GST-δLdPfn,.**(B)Western blots of purified LdPfn and δLdPfn, employing anti-LdPfn antibodies.** Mr, Molecular weight markers; Lane 1, GST-LdPfn; Lane 2, GST-δLdPfn. Asterisk and arrow head mark the positions of GST-LdPfn and GST-δLdPfn, respectively.

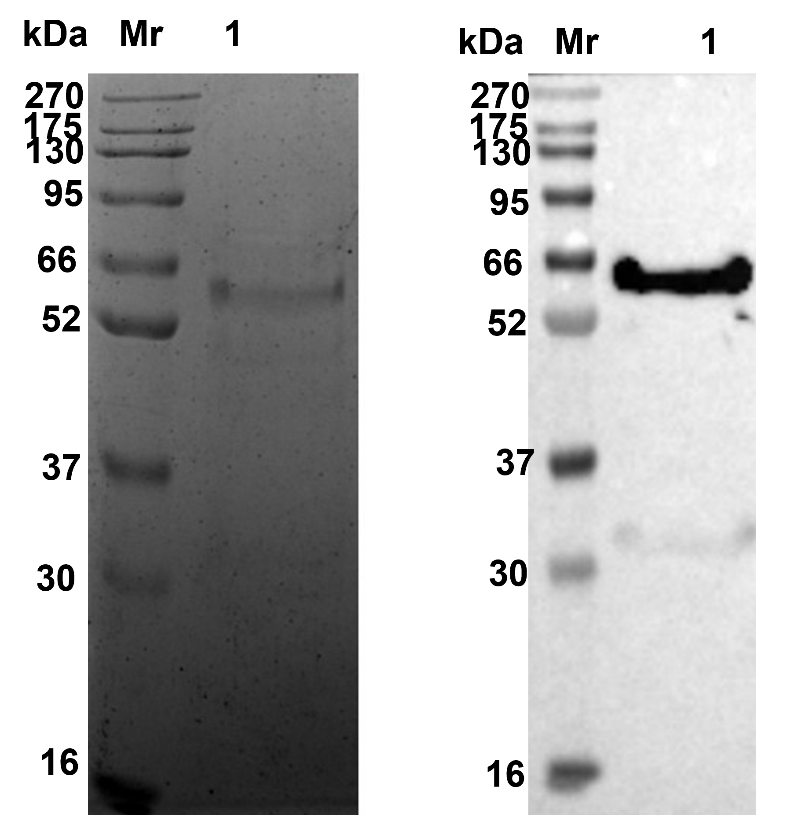

**S3Fig: (A) Coomassie blue stained 12% SDS-polyacrylamide gel showing purity of recombinant GST-LdPorin.** Mr, Molecular weight markers; Lane 1, GST-LdPorin. **(B) Western blot analysis of purified LdPorin, employing anti-GST antibodies.** Mr, Molecular weight markers; Lane 1, GST-LdPorin.
